# Ubiquitylome Analysis Reveals a Central Role for the Ubiquitin-Proteasome System in Plant Innate Immunity

**DOI:** 10.1101/2020.09.15.298521

**Authors:** Xiyu Ma, Chao Zhang, Do Young Kim, Yanyan Huang, Ping He, Richard D. Vierstra, Libo Shan

**Affiliations:** Department of Biochemistry and Biophysics, Texas A&M University, College Station, TX 77843, USA; Department of Plant Pathology and Microbiology, Texas A&M University, College Station, TX 77843, USA; Department of Genetics, 425-G Henry Mall, University of Wisconsin-Madison, Madison, WI 53706, USA; Advanced Bio Convergence Center, Pohang Technopark, Gyeong-Buk, South Korea, 37668; Department of Biology, Washington University in St. Louis, St. Louis, MO 63130, USA

**Keywords:** innate immunity, disease resistance, ubiquitylome, protein ubiquitylation, proteasome, mass spectrometry, Arabidopsis

## Abstract

Protein ubiquitylation profoundly expands proteome functionality and diversifies cellular signaling processes, with recent studies providing ample evidence for its importance to plant immunity. To gain a proteome-wide appreciation of ubiquitylome dynamics during immune recognition, we employed a two-step affinity enrichment protocol based on a 6His-tagged ubiquitin (Ub) variant coupled with high sensitivity mass spectrometry to identify *Arabidopsis* proteins rapidly ubiquitylated upon plant perception of the microbe-associated molecular pattern (MAMP) peptide flg22. The catalog from two-week-old seedlings treated for only 30 minutes with flg22 contained nearly 1,000 conjugates, 150 Ub footprints, and all seven types of Ub linkages, and included previously uncharacterized conjugates of immune components, such as RECEPTOR-LIKE KINASE 1 (RKL1) shown to negatively regulate plant immunity. *In vivo* ubiquitylation assays confirmed modification of several candidates upon immune elicitation, and revealed distinct modification patterns and dynamics for key immune components, including poly- and monoubiquitylation, as well as induced or reduced levels of ubiquitylation. Gene ontology and network analyses of the collection also uncovered rapid modification of the Ub-proteasome system itself, suggesting a critical auto-regulatory loop necessary for an effective MAMP-triggered immune response and subsequent disease resistance. Included targets were UBIQUITIN-CONJUGATING ENZYME 13 (UBC13) and proteasome component REGULATORY PARTICLE NON-ATPASE SUBUNIT 8b (RPN8b), whose subsequent biochemical and genetic analyses implied negative roles in immune elicitation. Collectively, our proteomic analyses further strengthened the connection between ubiquitylation and flg22-based immune signaling, identified novel components and pathways regulating plant immunity, and increased the database of ubiquitylated substrates in plants.

**One-sentence summary:** Proteome-wide catalogs of ubiquitylated proteins revealed a rapid engagement of the ubiquitin-proteasome system in Arabidopsis innate immunity.

## INTRODUCTION

Post-translational modifications (PTMs), such as phosphorylation, methylation, acetylation, glycosylation, myristoylation, and ubiquitylation diversify protein behaviors, thus increasing the functionality and regulation of the proteome. As one of the most abundant PTMs, ubiquitylation is involved in nearly all physiological and signaling processes in plants, including hormone perception, photomorphogenesis, circadian rhythms, self-incompatibility, and defense against biotic and abiotic challenges (Guerra and Callis, 2012; Trujillo and Shirasu, 2010; Vierstra, 2009; Zhou and Zeng, 2017). Attachment of the 76-amino acid ubiquitin (Ub) moiety occurs most commonly to accessible lysines (K) within the target via an isopeptide bond that links to the carboxyl (C)-terminal glycine of Ub.

This conjugation is typically mediated through a stepwise enzymatic cascade consisting of a Ub-activating enzyme (UBA, or E1), a Ub-conjugating enzyme (UBC, or E2), and finally, a Ub-protein ligase (or E3), the latter of which has greatly expanded and diversified in plants to recognize a plethora of substrates. Additional Ubs can become attached to one or more of the seven lysine residues (K6, K11, K27, K29, K31, K48, and K63) or the amino (N)-terminal amide nitrogen within previously attached Ubs, resulting in a myriad of poly-Ub chain architectures that provide additional functional diversity (Komander and Rape, 2012; Vierstra, 2009). The most common internal linkage involves lysine 48 (K48) whose poly-Ub chains mainly target proteins for breakdown by the 26S proteasome, a multisubunit proteolytic complex that recognizes the Ub moieties (Vierstra, 2009; Zhou and Zeng, 2017). Although not as well understood in plants (Paez Valencia et al., 2016; Romero-Barrios and Vert, 2018; Zhou and Zeng, 2017), other types of poly-Ub linkages, involving K6, K11, K27, K29, K33, or K63, and monoubiquitylation in which only a single Ub is attached, have also been connected to many intracellular events, including protein trafficking, signal transduction, protein-protein interactions, and protein degradation through autophagy (Komander and Rape, 2012).

In recent years, a myriad of plant defenses against a range of pathogens has become apparent. The front line of defense is to block pathogen entry by erecting physical barriers, such as wax layers, cell walls, cuticular lipids, cutin, and callose, which can be strengthened during infection (Thordal-Christensen, 2003). As a second line of defense, plants rely on innate immune responses to fend off infections internally, which are elicited upon recognition of pathogen- or microbe-associated molecular patterns (PAMPs/MAMPs) for pattern-triggered immunity (PTI), or pathogen-encoded effectors for effector-triggered immunity (ETI), which are encoded within the pathogen (Spoel and Dong, 2012; Zhou and Zhang, 2020). The host receptors known to detect these patterns/effectors are either receptor-like kinases (RLKs) or receptor-like proteins (RLPs) (Böhm et al., 2014; Couto and Zipfel, 2016). The best understood is the RLK receptor FLAGELLIN SENSING 2 (FLS2) and its co-receptor BRASSINOSTEROID INSENSITIVE 1-ASSOCIATED KINASE 1 (BAK1) that recognize the bacterial 22-amino-acid flagellin epitope - flg22. Subsequently, PTI signaling is relayed through receptor-like cytoplasmic kinases (RLCKs), the mitogen-activated protein kinase (MAPK) cascade, and transcription factors, that trigger diverse immune responses, including the production of reactive oxygen species (ROS) and the stress hormone ethylene (Couto and Zipfel, 2016; Yu et al., 2017). Pathogen effectors are recognized directly or indirectly by NUCLEOTIDE-BINDING SITE LEUCINE-RICH REPEAT (NBS-LRR) proteins (NLRs), which often leads to localized cell death at the infection sites known as the hypersensitive response (HR) (Cui et al., 2015). The defense hormone salicylic acid (SA) is indispensable for NLR-triggered HR (Cui et al., 2015).

Protein ubiquitylation is also emerging as a dynamic process in both PTI and ETI signaling (Adams and Spoel, 2018; Cheng and Li, 2012; Zhou and Zeng, 2017). As examples, the MAMP receptor FLS2 is ubiquitylated upon ligand engagement by the E3s PLANT U-BOX 12 (PUB12) and PUB13, resulting in FLS2 degradation (Lu et al., 2011; Zhou et al., 2015), while PUB13 also directs the ubiquitylation and degradation of LYSIN MOTIF RECEPTOR KINASE 5 (LYK5), an RLK that recognizes the fungal cell wall polysaccharide chitin (Liao et al., 2017). The RLCK *BOTRYTIS*-INDUCED KINASE 1 (BIK1), which provides a convergent signaling hub downstream of multiple MAMP receptors, is strongly regulated by Ub addition (Ma et al., 2020; Wang et al., 2018). Two U-box E3s, PUB25, and PUB26 polyubiquitylate BIK1 to control its steady-state levels, while the RING-type E3 ligases, RING-H2 FINGER A3A (RHA3A) and RHA3B, monoubiquitylate BIK1 to direct its endocytosis and signaling activation (Ma et al., 2020; Wang et al., 2018). Additionally, the PUB22, PUB23, and PUB24 E3s work collectively to negatively regulate PTI responses through modification of EXOCYST SUBUNIT EXO-70 FAMILY PROTEIN B2 (EXO70B2), which promotes vesicle trafficking (Stegmann et al., 2012). Levels of several NLR proteins, including RESISTANT TO *PSEUDOMONAS SYRINGAE* 2 (RPS2) and SUPPRESSOR OF *npr1-1* CONSTITUTIVE 1 (SNC1), are also regulated by CONSTITUTIVE EXPRESSOR OF PR GENES 1 (CPR1), which is the target-recognition F-Box subunit within an SKP1-CULLIN1-F-BOX (SCF) E3 complex (Cheng et al., 2011; Gou et al., 2012). Despite these examples, a systematic investigation of ubiquitylation during plant immune activation is currently lacking.

Recent advance in proteomics has enabled high-throughput and comprehensive views of ubiquitylomes under specific physiological conditions. The first catalog was generated with yeast in which wild-type Ub was replaced by a 6His-tagged variant to enable enrichment of ubiquitylated proteins by nickel-nitrotrilotriacetic acid (Ni-NTA) affinity chromatography under denaturing conditions, followed by liquid chromatography-mass spectrometric analysis (LC-MS) of the trypsinized preparations (Peng et al., 2003). Sarraco et al. (2009) then adopted this strategy for plants through the creation of a transgenic *Arabidopsis* line that constitutively expresses a hexa-6His-Ub concatamer which is post-translationally processed into functional 6His-Ub monomers. Subsequently, Ub-binding domains were exploited for the additional enrichment needed for more complex proteomes (Maor et al., 2007). Further improvements included Tandem-repeated Ub-Binding Entities (TUBEs) in which natural Ub-binding domains were oligomerized to increase enrichment and yield (Hjerpe et al., 2009). By combining TUBEs with the Ni-NTA affinity chromatography, stringent purifications were possible, which enabled the confident detection of over 1,000 ubiquitylated proteins in *Arabidopsis* (Aguilar-Hernandez et al., 2017; Kim et al., 2013). Notably, a number of Ub targets associated with disease resistance were identified.

Here, we employed this improved two-step TUBEs purification approach coupled with deep MS analysis of *Arabidopsis* seedlings before and shortly after exposure to flg22. The resulting ubiquitylome catalog revealed extensive proteome-wide changes soon after MAMP perception and uncovered a large number of ubiquitylated candidates potentially connected to plant innate immunity. We further observed by *in vivo* ubiquitylation assays distinct ubiquitylation patterns and dynamics for several immunity components, including key regulators that were modified by multiple ubiquitylation events. One target was the previously uncharacterized receptor protein kinase RECEPTOR-LIKE KINASE 1 (RKL1) that appears to negatively impact plant immunity. Interestingly, several Ub-proteasome pathway components were also confirmed as Ub targets upon PTI immune elicitation, including the E2 UBC13 and proteasome subunit REGULATORY PARTICLE NON-ATPASE SUBUNIT 8b (RPN8b). Taken together, our study identified novel connections between the Ub-proteasome system (UPS) and pathogen immunity as well as provided a valuable resource to further study this PTM during immune activation.

## RESULTS

### TUBEs enrichment of ubiquitylated *Arabidopsis* proteins upon flg22 treatment

To gain a global perspective of ubiquitylation during PTI elicitation, we employed a two-step TUBEs purification protocol to enrich for ubiquitylated proteins soon after exposure of two-week-old *A*. *thaliana* ecotype Col-0 seedlings to flg22 (Kim et al., 2013), which was based on a transgenic line harboring a synthetic *UBQ* gene encoding six 6His-Ub repeats expressed under the control of the Cauliflower mosaic virus (CaMV) *35S* promoter (Sarraco et al., 2009). The polymer is processed by endogenous deubiquitylases (DUBs) to release tagged and fully functional Ub monomers (Isono and Nagel, 2014). In the first step, extracts from *hexa-6His-UBQ* seedlings either untreated or treated for only 30 min to flg22 were prepared in a non-denaturing buffer containing the non-ionic detergent Triton X-100 and a protease inhibitor cocktail to minimize DUB activities that would disassemble Ub conjugates post homogenization, and then enriched for Ub-bearing species by TUBEs chromatography (Hjerpe et al., 2009; Raasi et al., 2004), using as the ligand four copies of the Ub-Associated Domain (UBA) sequence from human hHR23A interconnected through a short flexible linker, and fused with GLUTATHIONE S-TRANSFERASE (GST) (Figure 1A). Previous studies show that this GST-4X(UBA) fusion binds effectively to native Ub-conjugates bearing various chain topologies from *Arabidopsis*, thus allowing us to interrogate its near complete ubiquitylome (Aguilar-Hernandez et al., 2017; Kim et al., 2013). In the second step, we enriched for poly-His-containing proteins via Ni-NTA chromatography under strong denaturing conditions using buffers containing 7 M guanidine-HCl and 8 M urea for the application and washing steps, respectively, followed by elution with a buffer containing 8 M urea and 400 mM imidazole (Figure 1B).

**Figure 1.**
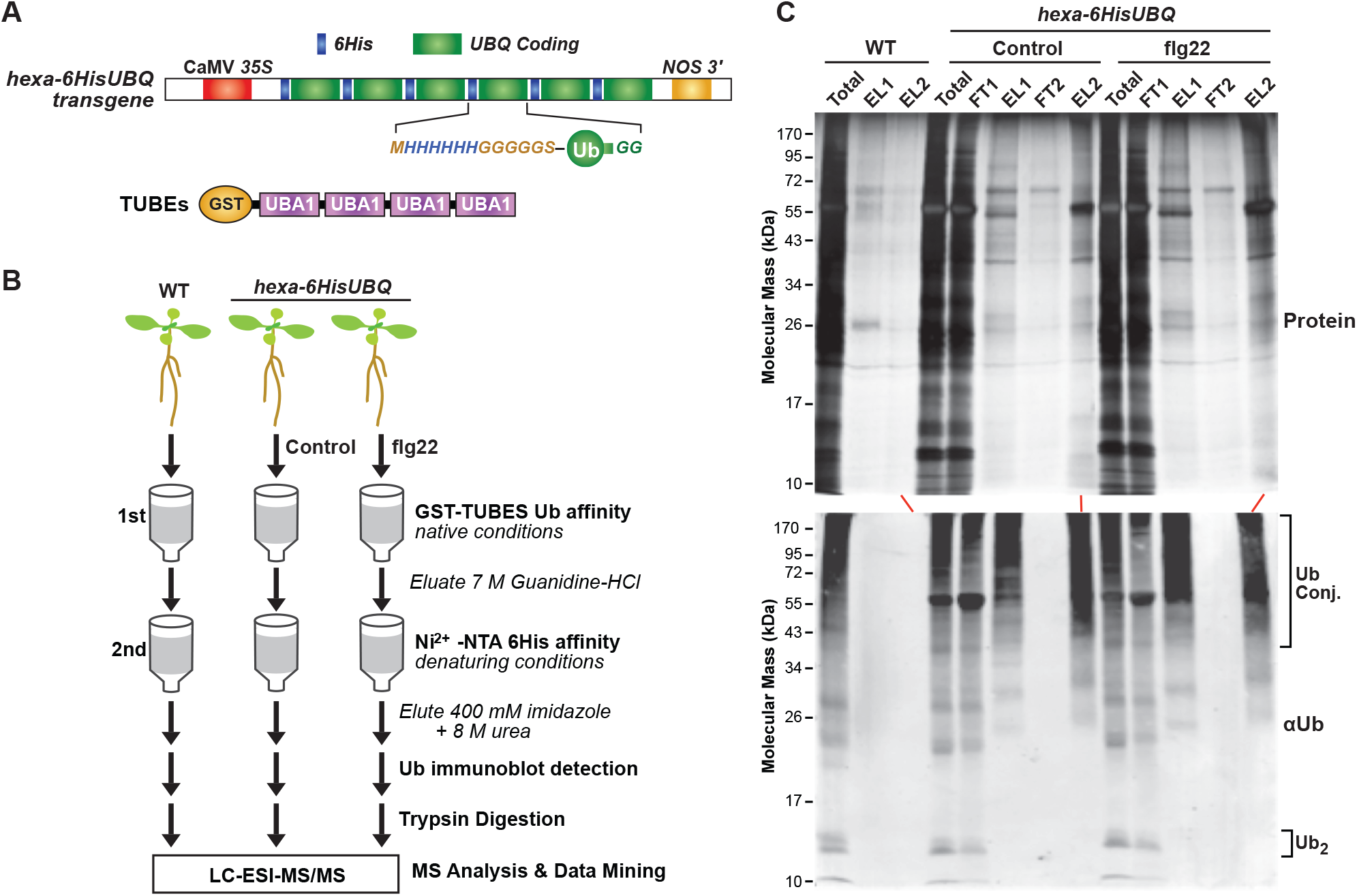
Two-step affinity purification of ubiquitylated proteins from transgenic *Arabidopsis* expressing *6HIS-UBQ* treated with or without flg22. **A.** Diagram of the *p35S:hexa-6HIS-UBQ* transgene*. A* synthetic *UBQ* gene encoding six repeats of a Ub monomer N-terminally tagged with a 6His sequence followed by a glycine (G)-rich linker (MHHHHHHGGGGGSA) was fused head-to-tail to form a single in-frame *hexa*-6*His-UBQ* transgene expressed under the control of the constitutive CaMV *35S* promoter (Saracco et al., 2009). **B.** Flow chart describing the proteomic approach to stringently identify Ub conjugates from *Arabidopsis p35S::hexa-6HIS-UBQ* seedlings. Ubiquitylated proteins were first enriched by the UBA-Ub affinity using GST-TUBEs beads under the native condition followed by Ni-NTA chromatography under the denaturing condition. The eluants were digested with trypsin and subjected to LC/ESI-MS/MS analysis. **C and D.** Ubiquitylated proteins isolated from *Arabidopsis* plants expressing 6His-Ub by the two-step affinity purification. Ubiquitylated proteins enriched as outlined in (**B**) were subjected to SDS-PAGE and by either stained for protein with silver (**C**) or immunoblotted with anti-Ub antibodies (**D**). Samples from wild-type seedlings (WT) purified with both GST-TUBEs and Ni-NTA beads were included for comparison. FT and EL represent the flow-through and eluted fractions, respectively, from the Ub affinity (1) and Ni-NTA affinity columns (2). The total represents the total protein extracts before affinity purifications. The migration positions of the Ub dimers (Ub_2_), and Ub conjugates (Ub-conj) are indicated by the brackets.

As seen in Figure 1C and D, eluants from the TUBEs and Ni-NTA columns prepared with control and flg22-treated *hexa-6His-UBQ* seedlings had substantially more ubiquitylated species than those obtained from wild-type seedlings, as detected by immunoblot analysis with rabbit anti-Ub antibodies. They appeared particularly enriched in high molecular mass Ub conjugates, with limited amounts of free Ub or the Ub dimer that were easily seen in total seedling extracts (Figure 1D). The main contaminant was RIBULOSE-1,5-BISPHOSPHATE CARBOXYLASE/OXYGENASE (Rubisco), which is often recognized by antibodies generated in rabbits.

### Identification of Ub-conjugates by LC-MS/MS

Ub-conjugates from the *hexa-6HIS-UBQ* plants were trypsinized, separated by reversed-phase nanoflow liquid chromatography (LC), and subjected to tandem MS (MS/MS) using a Velos LTQ mass spectrometer in the high energy collision mode (Figure 1B). Peptides were matched to the *Arabidopsis* ecotype Col-0 protein database (IPI database, version 3.85; http://www.arabidopsis.org) using SEQUEST, based on the identification of two or more different matching peptides with a ≤1% false discovery rate (FDR), or if a single matching peptide of equivalent stringency was identified that harbored a canonical Ub footprint (Kim et al., 2013). This footprint was detected by the presence of a di-Gly remnant (Gly-Gly) from Ub isopeptide linked to a lysine within the peptide, which increased the monoisotopic mass of the residue by 114 Da as well as protected the site from trypsin cleavage (Peng et al., 2003; Saracco et al., 2009). To help eliminate contaminants that bind non-specifically to the TUBEs and/or Ni-NTA columns, a background LC-MS/MS database was also generated with wild-type plants that underwent the same two-step affinity protocol; these proteins were subtracted from *hexa*-*6HIS-UBQ* datasets to obtain the final ubiquitylome catalog (Kim et al., 2013). In total, we generated MS/MS datasets from three biological replicates for control, untreated, and flg22-treated seedlings for final comparisons.

The summation of this MS analysis enabled the detection of 391 Ub conjugates in control samples (204, 210, 250 proteins for biological replicates 1, 2, 3, respectively) and 570 Ub conjugates in the flg22-treated samples (391, 400, 142 proteins for biological replicates 1, 2, 3, respectively) (Supplemental Table 1). Notably, 90 proteins in the control samples and 88 proteins in the flg22-treated samples were consistently identified in all three biological replicates, which likely represented a high abundance of ubiquitylated species. When combined, we uncovered 961 ubiquitylated proteins with high confidence; 690 were confined to either control or flg22-treated samples, whereas 271 proteins were present in both (Figure 2A). We detected 299 proteins unique to the flg22-treated samples versus 120 proteins unique to the control samples, implying that the PTI elicitation rapidly and markedly increases ubiquitylation in Arabidopsis (Figure 2A).

**Figure 2.**
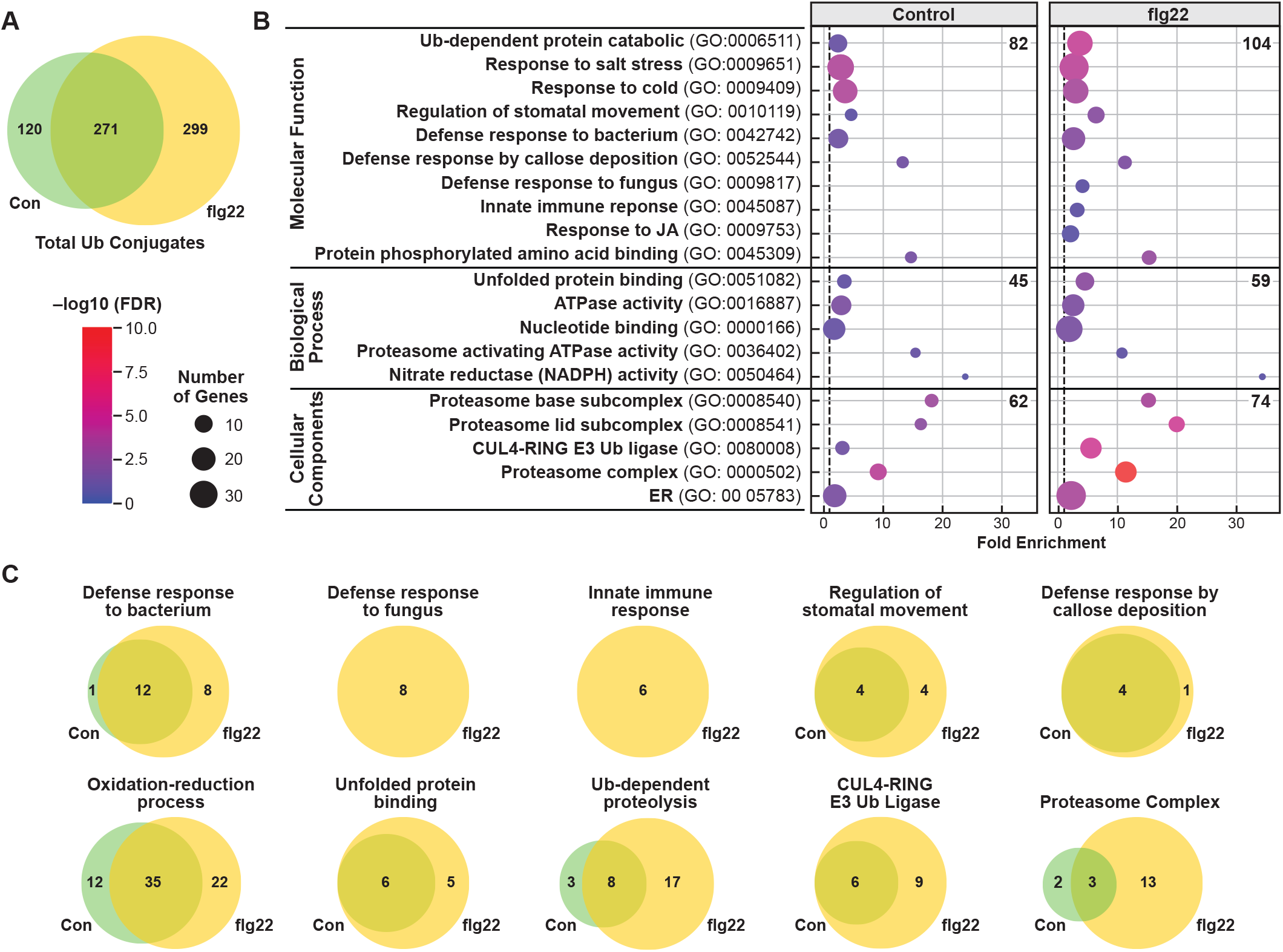
Gene ontology (GO) analysis of flg22-treated and untreated ubiquitylomes. **A.** Venn diagram showing the overlap of the total ubiquitylated proteins isolated from plants with or without flg22 exposure. **B.** Gene ontology (GO) enrichment of ubiquitylated proteins in control or flg22 treated ubiquitylomes. GO analysis in biological processes, molecular functions, and cellular components was performed using the DAVID Functional Annotation Tool (Huang da et al., 2009) and GO annotations from the TAIR GO database (http://www.arabidopsis.org/). The fold enrichment was calculated based on the frequency of proteins annotated to the term compared with their frequency in the total proteome. The dot size indicates the number of genes associated with each process and the dot color indicates the significance of the enrichment. FDR was based on the corrected *p*-values. The vertical grey dashed line represents a fold enrichment of 1. The total number of GO terms was listed under each GO domain (biological processes, molecular functions, and cellular components). The selected terms related to stress and the UPS are shown. **C.** Venn diagram showing the overlaps of ubiquitylated proteins with or without flg22 in the selected categories. Proteins were categorized into functional groups based on the GO annotations; those related to defense responses and the UPs are shown.

### Gene ontology analyses of flg22-treated and -untreated ubiquitylomes

To gain a global perspective for how ubiquitylation is influenced by immune elicitation, we classified the identified targets in the control and flg22-treated ubiquitylomes based on their known or predicted functions within the Gene Ontology (GO) database using the DAVID Functional Annotation Tool (Huang da et al., 2009). As shown in Figure 2B, the total number of significantly enriched GO terms (FDR <0.05) was larger for the flg22-treated ubiquitylome than for that for the control for the three main GO domains (‘Biological Processes’: 104 vs. 82, ‘Molecular Functions’: 59 vs. 45, and ‘Cellular Components’ 74 vs. 62), and were noticeably more dedicated to pathogen-related events. Enrichments included the GO terms ‘defense response to fungus’, and ‘innate immune response’, which were found only in the flg22-treated ubiquitylome, while the related terms ‘defense response to bacterium’, ‘regulation of stomatal movement’, ‘defense response by callose deposition’, and ‘oxidation-reduction process’ were more significantly enriched (Figure 2B, C).

As compared to the total proteome, we also saw significant enrichment for GO terms associated with ‘Ub-dependent protein catabolic process’ (Biological process), and ‘Cul4-RING E3 Ub ligase’ (Cellular components) in both control and flg22-treated ubiquitylomes, as were the GO terms related to stress, such as ‘response to salt stress’ and ‘response to cold stress’ (Figure 2B). Strikingly, this enrichment in Ub-related events was more robust in the flg22-treated versus control ubiquitylomes (3.7-fold vs. 2.4-fold for ‘ubiquitin-dependent protein catabolic process’ and 5.5-fold vs. 3.2-fold for ‘Cul4-RING E3 Ub ligase’, respectively). A prevalence for UPS components in the flg22-treated samples was also evident for proteins associated with ‘proteasomes’ (11.4-fold vs. 9.2-fold, respectively; Figure 2C). Taken together, this non-targeted MS analysis directly connected rapid flg22-induced ubiquitylation to Arabidopsis proteins associated with immunity and the UPS itself.

### Global ubiquitylation of immunity-related proteins

Among the identified Ub conjugates were many key immunity-related proteins (Table 1). To link these ubiquitylated species to immune signaling globally, we generated an interaction network using the Search Tool for Retrieval of Interacting Genes/Proteins (STRING) database (Szklarczyk et al., 2015), which revealed an interconnected web with defense-related proteins often situated at key hubs (Figure 3A, B). Examples included plasma membrane-resident leucine-rich repeat-RLKs (LRR-RLKs), such as RECEPTOR-LIKE KINASE 7 (RLK7) and SUPPRESSOR OF *BIR1-1*/EVERSHED (SOBIR1/EVR) that interact with multiple LRR-RLP receptors (Figure 3A, B). Whereas SOBIR1 serves as a common component in RLP-mediated plant immunity (Albert et al., 2015; Liebrand et al., 2014), RLK7 is a receptor for PAMP-INDUCED PEPTIDE 1 (PIP1), which acts as a damage-associated molecular pattern (DAMP) to amplify plant immune signaling (Hou et al., 2014). Additionally, LysM RLK1-INTERACTING KINASE 1 (LIK1), which was specific to the flg22-treated ubiquitylome, has been shown to interact with the fungal chitin receptor complex component CERK1 to positively regulate resistance against necrotrophic pathogens (Le et al., 2014).

**Figure 3.**
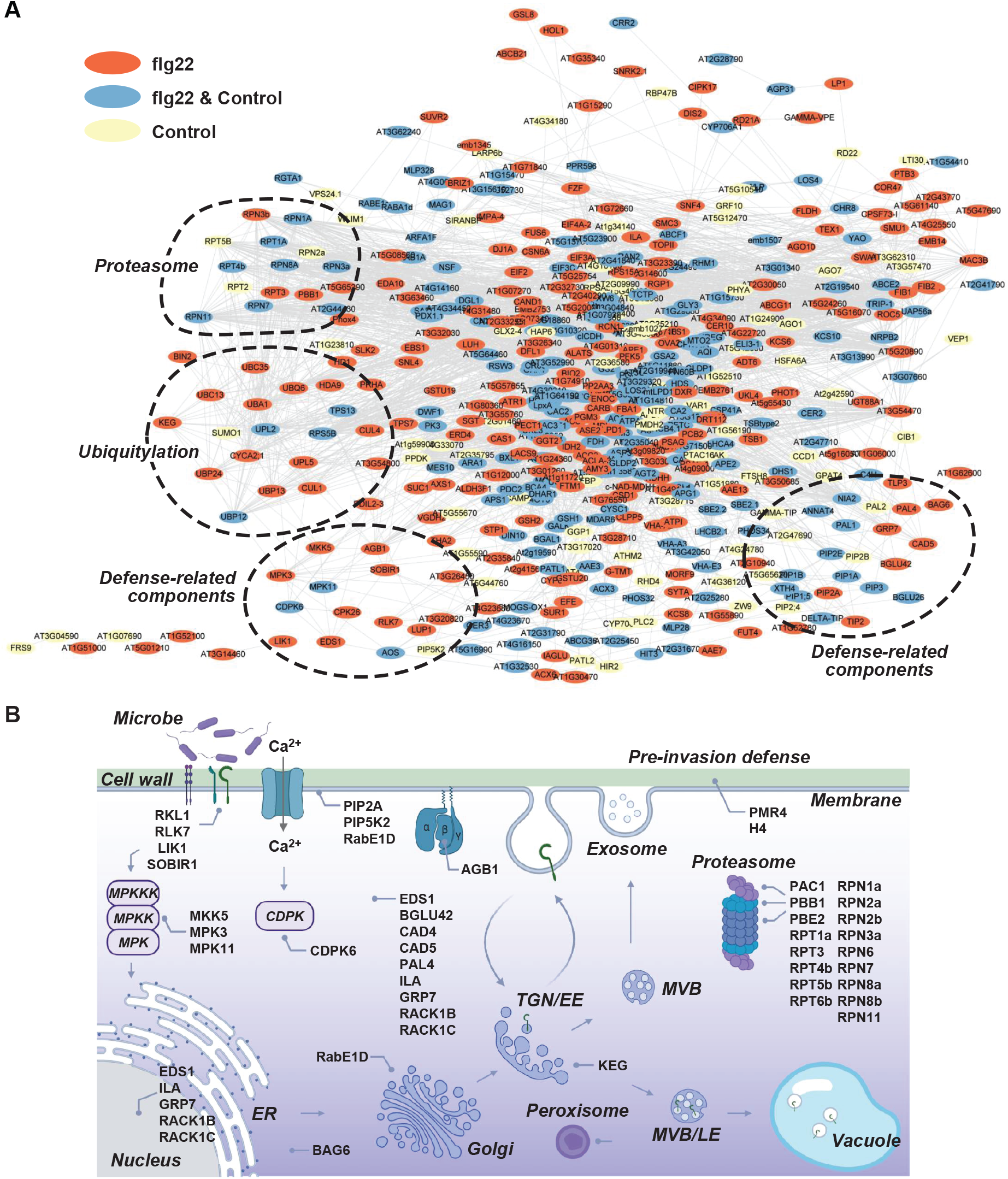
Protein interaction network of ubiquitylated proteins identified from 6*HIS-UBQ* seedlings treated with or without flg22. **A.** Protein interaction networks were generated with the list of 391 (control) and 570 (flg22-treated) ubiquitylated proteins using the STRING database and visualized in Cytoscape. The node colors indicate proteins identified in control (yellow), flg22-treated (red), or both conditions (blue) of datasets. Clusters of the proteasome, ubiquitylation, and defense-related components are marked by dashed lines. **B.** Map of proposed subcellular locations and functions of immunity-related proteins identified from the ubiquitylomes.

**Table 1.** Ubiquitylation of immunity-related proteins identified from *6HIS-UBQ* seedlings treated with or without flg22.

| Locus ID | Other names | Description | Category | Role in immunity | Condition | Ref. |
| --- | --- | --- | --- | --- | --- | --- |
| AT4G34230 | CAD5 | Cinnamyl alcohol dehydrogenase 5 | Preformed defense | Involves in lignin biosynthesis and disease resistance against *Pst* DC3000. | flg22 | (Tronchet et al., 2010) |
| AT3G10340 | PAL4 | Phenylalanine ammonia-lyase 4 | Preformed defense | Functions in the lignin and SA pathway in plant disease resistance. | flg22 | (Chen et al., 2017; Huang et al., 2010) |
| AT5G36890 | BGLU42 | Beta glucosidase 42 | Preformed defense | Promotes disease resistance against *B.cinerea*, *H.arabidopsidis* and *Pst* DC3000 | flg22 | (Stringlis et al., 2018; Zamioudis et al., 2014) |
| AT3G53420 | PIP2A, PIP2;1 | Plasma membrane intrinsic protein 2A | Preformed defense | Involves in stomatal closure triggered by abscisic acid (ABA) or flg22 for water and H<sub>2</sub>O<sub>2</sub> transport activities | flg22 | (Rodrigues et al., 2017) |
| AT1G09970 | RLK7 | Leucine-rich repeat receptor-like kinase 7 | RLK | Receptor for PIP1 defense peptide in PTI signaling | flg22 | (Hou et al., 2014; Pitorre et al., 2010) |
| AT3G14840 | LIK1 | LysM RLK1-interacting kinase 1 | RLK | Interacts with CERK1 and positively regulates resistance to necrotrophic pathogens. | flg22 | (Le et al., 2014) |
| AT1G48480 | RKL1 | Receptor-like protein kinase 1 | RLK | Characterized in this study | flg22 | (Tarutani et al., 2004) |
| AT2G31880 | SOBIR1, | Leucine-rich repeat | RLK | Interacts with multiple | flg22 | (Albert et |
|  | EVR | receptor-like protein |  | LRR-RLP and functions as a common component in RLP-mediated plant immunity |  | al., 2015; Liebrand et al., 2014) |
| AT1G53350 | RPP8L2 | RPP8 like protein 2; Resistance protein; CC-NBS-LRR | CNL | Belongs to CNL-D subclass, may involve in controlling virus infection | flg22 | (Bakker et al., 2006; Mondragon-Palomino et al., 2017; Revers et al., 2003; Tan et al., 2007) |
| AT5G45240 |  | Resistance protein; TIR-NBS-LRR | TNL | uncharacterized | flg22 |  |
| AT5G48770 |  | Resistance protein; TIR-NBS-LRR | TNL | uncharacterized | flg22 |  |
| AT5G13530 | KEG | RING E3 Ub-protein ligase; KEEP ON GOING | Signaling | Essential for the secretion of apoplast antimicrobial proteins. | flg22 | (Gu and Innes, 2011) |
| AT2G46240 | BAG6 | BCL-2-associated athanogene 6 | Signaling | Co-chaperone regulating diverse cellular pathways, such as programmed cell death, abiotic stress responses, and plant basal resistance | flg22 | (Li et al., 2016) |
| AT2G21660 | CCR2, GRP7 | Cold, circadian rhythm, and RNA binding 2; glycine-rich RNA binding protein 7 | Signaling | Associates with FLS2 mRNA and proteins; maintains the FLS2 protein level | flg22 | (Nicaise et al., 2013) |
| AT4G34460 | AGB1,<br>ELK4 | Heterotrimeric G-protein beta subunit; ERECTA-LIKE 4 | Signaling | Complexes with FLS2 and regulates BIK1 stability | flg22 | (Liang et al., 2016; Liu et al., 2013) |
| AT5G03520 | RAB8C,<br>RABE1D | Rab GTPase homolog 8C | Signaling | Involves in protein trafficking, interacts with AvrPto, and accumulates in response to bacterial infection | flg22 | (Hou and Gao, 2017; Speth et al., 2009a) |
| AT3G21220 | MEK5,<br>MKK5 | Mitogen-activated protein kinase kinase 5 | Signaling | Activate MPK3 and MPK6 in PTI signaling. | flg22 | (Meng and Zhang, 2013) |
| AT3G45640 | MPK3 | Mitogen-activated protein kinase 3 | Signaling | Activated by MAMP treatment in PTI signaling | flg22 | (Asai et al., 2002; Pecher et al., 2014; Su et al., 2018) |
| AT2G33340 | MAC3B;<br>PUB60 | MOS4-associated complex 3B; Plant U-Box 60 E3 ligase | Signaling | Localizes to the nucleus and involves in plant innate immunity. | flg22 | (Monaghan et al., 2009) |
| AT1G77740 | PIP5K2 | Phosphatidylinositol-4-phosphate 5-kinase | Signaling | Responsible for PI(4,5)P2 biosynthesis; inhibits fungal pathogen development and triggers disease resistance | Control | (Qin et al., 2020) |
| AT3G18130 | RACK1C | Receptor for activated C kinase 1C | Signaling | scaffold proteins connecting heterotrimeric G proteins with a MAPK cascade in plant immunity | Control, flg22 | (Cheng et al., 2015; Nakashima et al., 2008; Su et al., 2015) |
| AT3G48090 | EDS1 | Enhanced disease susceptibility 1; a lipase-like protein | Signaling | Required for multiple TNL-mediated resistance. | flg22 | (Cui et al., 2015; Heidrich et al., 2011; Parker et al., 1996) |
| AT3G50480 | HR4 | Homolog of RPW8 | Signaling | Involved in autoimmunity mediated by RPP7 oligomerization | flg22 | (Li et al., 2020) |
| AT1G64790 | ILA | ILITYHIA; HEAT repeat protein | SAR | Involved in immunity against bacterial infection, non-host resistance, and systemic acquired resistance | flg22 | (Monaghan and Li, 2010) |
| AT2G44490 | BGLU26, PEN2 | Beta glucosidase 26, PENETRATION 2 | Preformed defense | Required for callose deposition and glucosinolate activation in pathogen-induced resistance | Control, flg22 | (Clay et al., 2009; Pastorczyk and Bednarek, 2016) |
| AT3G19450 | CAD4 | Cinnamyl alcohol dehydrogenase 4 | Preformed defense | Involved in lignin biosynthesis and disease resistance to bacterial pathogens. | Control, flg22 | (Kim et al., 2004; Tronchet et al., 2010) |
| AT4G03550 | GSL5, PMR4 | Callose synthase | Preformed defense | Required for callose formation induced by pathogen infections. | Control, flg22 | (Jacobs et al., 2003; Wawrzynska et al., 2010) |
| AT1G63350 |  | Resistance protein; CC-NBS-LRR | CNL | uncharacterized | Control, flg22 |  |
| AT1G01560 | MPK11 | MAP Kinase 11 | Signaling | Activated upon flg22 treatment | Control, flg22 | (Bethke et al., 2012; Eschen-Lippold et al., 2012) |
| AT4G23650 | CPK3 | Calcium-dependent protein kinase 3 | Signaling | required for sphingolipid-induced cell death and viral infection | Control, flg22 | (Lachaud et al., 2013; Perraki et al., 2017) |
| AT1G48630 | RACK1B | Receptor for activated C kinase 1B | Signaling | scaffold proteins connecting heterotrimeric G proteins with a MAPK cascade in plant immunity | Control, flg22 | (Cheng et al., 2015; Nakashima et al., 2008; Su et al., 2015) |

Downstream of the immune receptor complexes are sequential phosphorylation events involving MAPK cascades driven by MAPK KINASE KINASES (MAPKK/MEKK), MAPK KINASES (MAPKK/MKK), and MAPKS (MPK) components (Meng and Zhang, 2013). Notably, MKK5, MPK3, and MPK11, which become phosphorylated upon flg22 perception, were detected here, with MKK5 and MPK3 only identified in the flg22-treated ubiquitylome (Figure 3B). Heterotrimeric G-proteins, consisting of Gα, Gβ, and Gγ subunits, act as molecular switches to mediate diverse signal transduction events (Urano et al., 2013). In particular, *ARABIDOPSIS* G PROTEIN β-SUBUNIT1 (AGB1), which associates with FLS2 and regulates BIK1 stability during plant PTI (Liang et al., 2016; Liu et al., 2013), was identified in the flg22-treated samples. Ubiquitylated forms of RECEPTOR FOR ACTIVATED C KINASE (RACK)-1B and RACK1C, which scaffolds and connects heterotrimeric G proteins with MAPK cascades to form a unique signaling pathway in plant immunity (Cheng et al., 2015), were identified in both control and flg22-treated ubiquitylomes, or only the flg22-treated ubiquitylome, respectively.

Protein trafficking, regulated by a cascade of factors such as small GTPases from Rab subfamilies, also plays important roles in plant immunity (Gu et al., 2017). RabE1D, identified here in the flg22-treated ubiquitylome, is associated with Golgi and the plasma membrane and might regulate the apoplast secretion of anti-microbial proteins, such as PATHOGENESIS-RELATED PROTEIN 1 (PR1) that induce resistance against *Pseudomonas syringae* pv. *tomato* (*Pst*) DC3000 (Speth et al., 2009b). Likewise, the E3 KEEP ON GOING (KEG), which localizes to the trans-Golgi network/early endosome (TGN/EE) compartments and is essential for TGN/EE-mediated secretion of PR1 and papain-like Cys protease C14 (Gu and Innes, 2011) appears to be ubiquitylated by a flg22-induced event (Figure 3B).

ETI components were also identified as flg22-specific ubiquitylation substrates. Included were four uncharacterized NLR proteins (AT1G53350, AT1G63350, AT5G45240, and AT5G48770) along with ENHANCED DISEASE RESISTANCE 1 (EDS1), which is a nucleocytoplasmic lipase-like protein indispensable for multiple TOLL-INTERLEUKIN 1 RECEPTOR (TIR) domain NLR-mediated resistance events (Cui et al., 2015). Proteins implicated in long-distance resistance were identified in the flg22-treated ubiquitylome, such as the HEAT-repeat protein ILITHYIA (ILA) that encourages systemic acquired resistance (Monaghan et al., 2010). The peroxisome-localized glycol hydrolase PENETRATION2 (PEN2) was seen in all control and flg22-treated samples and thus considered to be a high abundance Ub target. PEN2, together with PEN1 and PEN3, actuate a cell wall-based defense against non-adapted pathogens by limiting pathogen entry (Figure 3B) (Clay et al., 2009; Collins et al., 2003; Fuchs et al., 2016). Similarly, the lignin biosynthetic enzymes, CINNAMYL ALCOHOL DEHYDROGENASE 4 (CAD4), identified in both control and flg22-treated ubiquitylomes, and CAD5 identified only in flg22-treated ubiquitylomes, likely protect against pathogen invasion by modifying the lignin content of cell walls (Tronchet et al., 2010). The callose synthase POWDERY MILDEW RESISTANT 4 (PMR4), which is required for wound- and/or pathogen-induced callose formation (Meyer et al., 2009; Wawrzynska et al., 2010), was also found in both datasets (Figure 3B). Taken together, our ubiquitylome profiles implicate Ub addition in multiple defense responses ranging from the preformed cell wall-based protection, to inducible PTI and ETI, and systemic defense (Figure 3B).

### Distinct ubiquitylation patterns and dynamics of immunity-related proteins upon flg22 elicitation

To examine more closely the ubiquitylation dynamics (+/−flg22) of several immunity-related substrates, we exploited a previously established *in vivo* ubiquitylation system in which FLAG-tagged Ub (FLAG-Ub) is co-expressed with HA- or myc-tagged targets in wild-type Arabidopsis leaf protoplasts (Zhou et al., 2014). Ubiquitylated proteins are enriched by immunoprecipitation with anti-FLAG antibody beads and then probed by immunoblot analysis with anti-HA or anti-myc antibodies to assay for possible ubiquitylated species.

As shown in Figure 4A, we successfully confirmed the ubiquitylation of MPK3, MKK5, SOBIR1, and EDS1 using this *in vivo* system combined with HA or myc-tagged versions. For MPK3-HA, multiple ubiquitylated species were evident in anti-FLAG immunoprecipitates with a dominant form seen at ∼52 kDa. Being ∼8 kDa bigger than that of unmodified MPK3-HA (∼43.8 kDa), we consider it likely that it represents a monoubiquitylated form. Notably, ubiquitylation of MPK3-HA was transiently enhanced soon after flg22 treatment (10 min) but quickly returned to a basal level of modification at 30 min (Figure 4A), even though the levels of unmodified MPK3-HA remained steady for one hr of flg22 treatment (Figure 4A), suggesting that MPK3 ubiquitylation triggered by flg22 helps relay the flg22 signal. By contrast, modification of MKK5-myc, easily seen by a strong monoubiquitylated species, appeared to be constitutive, *i.e*., its level was unchanged by flg22 elicitation (Figure 4B). For both SOBIR1-HA and EDS1-HA, poly-ubiquitylated species accumulated in the *in vivo* system. While, the smear of high molecular mass conjugates for SOBIR1 was strongly enhanced upon flg22 treatment, they were diminished for EDS1 (Figure 4C and 4D). Taken together, we confirmed the ubiquitylation of candidate proteins using this cell-based assay, which revealed surprisingly distinct ubiquitylation patterns and dynamics for the candidates upon immune elicitation with flg22.

**Figure 4.**
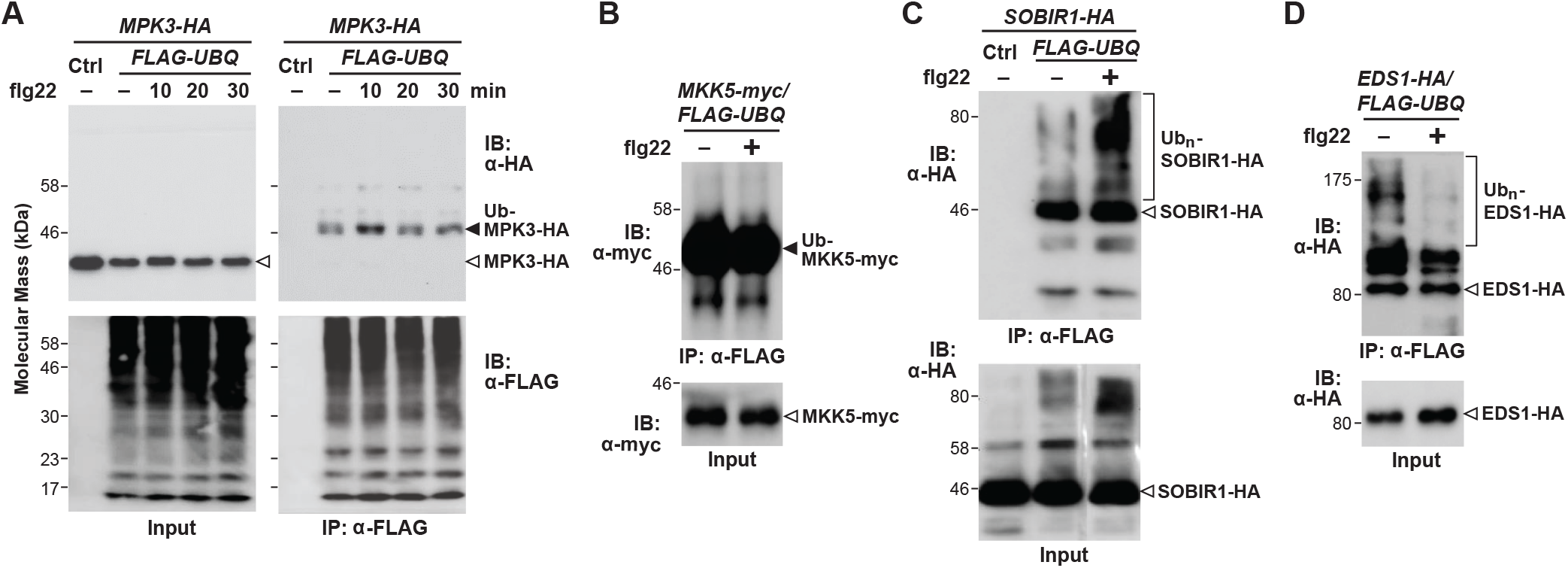
*In vivo* confirmation that several candidates are ubiquitylated in *Arabidopsis*. *Arabidopsis* wild-type protoplasts were co-transfected with FLAG-tagged Ub (*FLAG-UBQ*) and HA- or myc-tagged substrate or a control vector (Ctrl) and incubated for 10 hr followed by treatment with 100 nM flg22 for the indicated. Protein extracts were immunoprecipitated with anti-FLAG agarose beads (IP: α-FLAG) and the ubiquitylated proteins were immunoblotted with anti-HA or anti-myc antibodies (top left) or anti-FLAG antibodies (bottom left). The input controls were shown by an anti-HA/myc (top right) or anti-FLAG (bottom right) immunoblots. **A.** Monoubiquitylation of MPK3 is enhanced upon flg22 treatment. Protoplasts were isolated at 0 to 30 min after flg22 exposure. **B.** Polyubiquitylation of SOBIR1 *in vivo* is enhanced upon a 30 min treatment with flg22. **C, D.** MKK5, and EDS1 were ubiquitylated *in viv*o following a 30-min treatment with flg22. The above experiments were performed three times with similar results.

### Ubiquitylation of RKL1 and its role in plant immunity

The LRR III protein RECEPTOR-LIKE KINASE 1 (RKL1), identified here as a flg22-triggered ubiquitylation substrate, was previously shown to modulate the activity of the aquaporin PLASMA MEMBRANE INTRINSIC PROTEIN (PIP) family during osmotic and oxidative stress together with its close homolog RLK902 (Bellati et al., 2016). Intriguingly, while RKL1 was not yet connected to pathogen defense, RLK902 had been shown to transmit an immune signal by phosphorylating the RLCK BRASSINOSTEROID-SIGNALING KINASE 1 (BSK1) (Zhao et al., 2019). The RKL1 protein harbors a five-LEUCINE RICH REPEAT (LRR) extracellular domain, a transmembrane domain, and a cytosolic kinase domain (Figure 5A), and is mainly expressed in the stomata, hydathodes, and trichomes of young rosette leaves, and floral organ abscission zones based on transcriptomic analyses (Tarutani et al., 2004). When expressed in our leaf protoplast system with FLAG-Ub, we found that RKL1-HA was constitutively poly-ubiquitylated *in vivo*, despite our proteomic analyses of seedlings which found these species only in flg22-treated samples (Figure 5B).

To further characterize the role(s) of RKL1 in PTI, we assayed both MAMP-induced ROS production and bacteria disease spread, using a T-DNA insertion mutant compromising RKL1 function available from the SALK insertion collection (Alonso et al., 2003). The *rkl1-1* mutant (*SALK_099094*) bears an insertion within the first exon (Tarutani et al., 2004), which was confirmed here by genomic PCR analysis using gene-specific and T-DNA border primers (Figure S1A). When compared to wild-type *Arabidopsis* Col-0 leaves, we found that homozygous *rkl1-1* leaves displayed enhanced ROS production upon flg22 treatment based on a chemiluminescence assay quantifying luminol oxidation. We also conducted an assay for disease spread by infiltrating *Arabidopsis* leaves with the bacterial pathogen *Pst* DC3000 and then measuring bacterial numbers 2 days afterwards. As shown in Figure 5D, the *rkl1-1* mutant displayed significantly less infection as compared to wild type, consistent with a robust defense (Figure 5D). Together, the data implicated for the first time that RKL1 is a negative regulator of plant immunity, which is possibly influenced by ubiquitylation.

**Figure 5.**
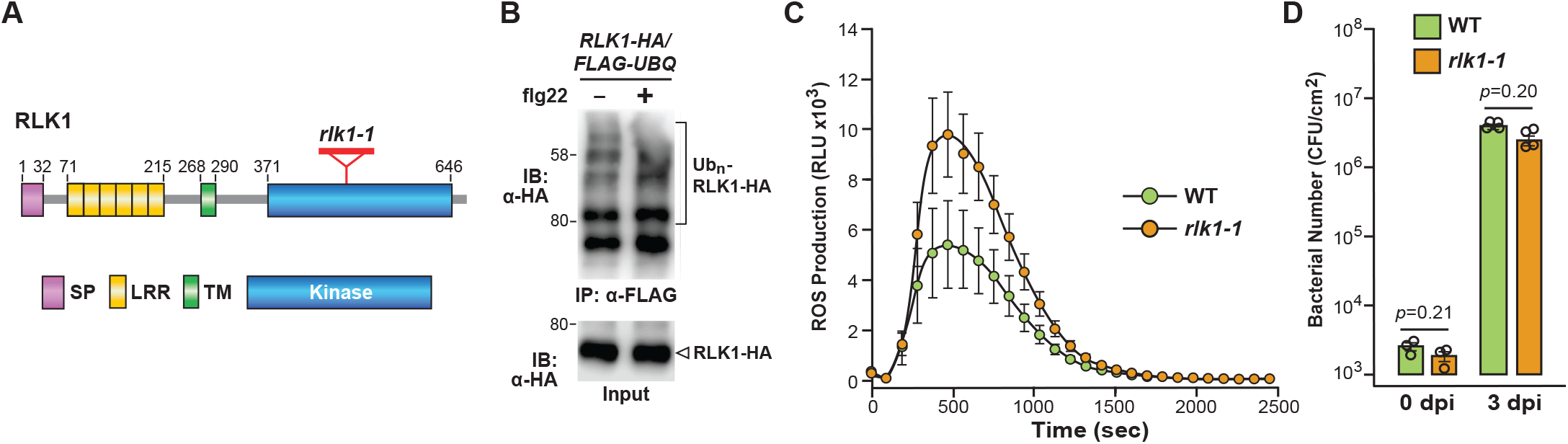
RKL1 is ubiquitylated and involved in PTI responses. **A.** Diagram of the RKL1 protein. The signal peptide (SP), leucine-rich repeat (LRR), transmembrane (TM), and protein kinase (K) domains are shown. The amino acids demarcating each domain are shown. **B.** *In vivo* ubiquitylation of RKL1 upon flg22 treatment. Protoplasts were co-transfected with *FLAG-UBQ* and HA-tagged RLK1 (*RKL1-HA*) or a control vector (Ctrl) and incubated for 10 hr followed by treatment with 100 nM flg22 for 30 min. Proteins were immunoprecipitated with anti-FLAG agarose beads (IP: α-FLAG) and the ubiquitylated proteins were detected by immunoblotting with anti-HA antibodies (top). The input controls were shown by an anti-HA immunoblot (bottom). **C.** flg22-induced reactive oxygen species (ROS) burst is elevated in the *rkl1-1* mutants. Leaf disks of four-week-old soil-grown *Arabidopsis* WT or *rkl1-1* plants (*SALK 099094*) were treated with 100 nM flg22 or water as a control, and ROS production was monitored over time. The data are shown as mean ± s.e.m (n = 16) of relative fluorescence units (RLU) overlaid on the dot plot. **D.** *rkl1* mutants have elevated resistance against *Pst* DC3000. Leaf disks of four-week-old soil-grown *Arabidopsis* WT or *rkl1-1* plants were hand-infiltrated with *Pst* DC3000 at OD_600_=5×10^−4^ CFU/ml, and bacterial growth was counted at 0 or 3 dpi. The data are shown as mean ± s.e.m. overlaid on dot plot (n = 3 for 0 dpi; n = 4 for 3 dpi). The experiments were performed three times with similar results.

### Ubiquitylation of UPS components

As revealed above (Table 2; Figures 2 and 3), multiple UPS components were preferentially identified in flg22-treated ubiquitylome. Included were the E1 UBA1, the E2 isoforms UBC13, and UBC35 (UBC13A), the E3 components CULLIN 1 (CUL1), CUL4, and UB-PROTEIN LIGASE 5 (UPL5), components of the 26S proteasome, and the DUBs Ub-Specific Proteases (UBP)-13 and UBP24 and a predicted Ub-Carboxy-Terminal Hydrolase (UCH) (At4g24320) expected to release free Ub monomers from poly-Ub chains and poly-Ub translation products. UPL5 is part of the single polypeptide HECT E3 family. By contrast, CUL1, together with S-PHASE KINASE-ASSOCIATED PROTEIN 1 (SKP1), RING BOX 1 (RBX1), and one of possibly >700 F-box protein variants, assemble into SCF-type E3 complexes (Vierstra, 2009), while the CUL4 scaffolds DNA DAMAGE-BINDING (DDB)-type E3 complexes assembled with RBX1, and one of over 85 DWD box-containing DDB proteins (Vierstra, 2009). Previously, DDB1 was implicated in *PR* gene expression and resistance to *Agrobacterial* infection in tomato (Liu et al., 2012b), which might explain the connection between CUL4 and immune elicitation, while UPL5 is required for SA- and NONEXPRESSER OF *PR* GENES 1 (NPR1)-mediated plant immunity (Furniss et al., 2018c). Our discovery of the DUBs UBP13, UBP24, and a UCH in the flg22-treated ubiquitylome implies that the release of Ub from targets is also a regulatory step in immune elicitation.

**Table 2.** Examples of UBQ pathway components identified from ubiquitylomes

| Locus ID | Other names | Description | Category | Role in immunity | Condition | Ref. |
| --- | --- | --- | --- | --- | --- | --- |
| AT2G47110 | UBQ6 | Poly-UBQ gene | Ubiquitin | Not known. | flg22 |  |
| AT2G30110 | UBA1 | Ub-activating enzyme 1 | E1 | Involved in PTI and ETI. | flg22 | (Furniss et al., 2018b; Goritschnig et al., 2007) |
| AT1G78870 | UBC13A, UBC35 | Ub-conjugating enzyme 35/13A | E2 | Regulates the immune response mediated by Fen and other R proteins through Lys-63–linked ubiquitylation | flg22 | (Mural et al., 2013; Wang et al., 2019) |
| AT3G46460 | UBC13 | Ub-conjugating enzyme 13 |  | Characterized in this study. | flg22 |  |
| AT4G02570 | AXR6, CUL1 | Cullin-1, a component of SCF Ub E3 ligase complexes |  | SKP1-CULLIN1-CPR1 (SCF), regulates protein stability of SNC1 and RPS2 and autoimmunity | flg22 | (Cheng et al., 2011) |
| AT5G46210 | CUL4 | CULLIN4, Ub E3 ligase |  | Required for *PR* gene expression and resistance to Agrobacterial infection. | flg22 | (Liu et al., 2012a) |
| AT4G12570 | UPL5 | Ub E3 ligase UPL5 | E3 | Required for SA- and NPR1-mediated plant immunity | flg22 | (Furniss et al., 2018a; Miao and Zentgraf, 2010; Zhou and Zeng, |
|  |  |  |  |  |  | 2017) |
| AT1G70320 | UPL2 | Ub E3 ligase 2 |  | Not known. | Control, flg22 |  |
| AT4G30890 | UBP24 | Ub-specific protease 24 | UBP | Negatively regulates abscisic acid signaling; a potential MPK3 interactor. | flg22 | (Rayapuram et al., 2018; Zhao et al., 2016) |
| AT3G11910 | UBP13 | Ub-specific protease 13 |  | Negatively regulates plant resistance against *Pst* DC3000 infections | flg22 | (Ewan et al., 2011) |
| AT5G06600 | UBP12 | Ub-specific protease 12 |  |  | Control, flg22 |  |
| AT4G24320 |  | Ub carboxyl-terminal hydrolase family protein | UCH | Not known. | flg22 |  |
| AT3G61140 | CSN1, FUS6 | COP9 signalosome subunit 1 | COP9 | Not known. | flg22 |  |
| AT5G56280 | CSN6A | COP9 signalosome complex subunit 6a |  | Not known. | flg22 |  |
| AT3G22110 | PAC1 | 26S proteasome subunit alpha type-4 | Proteasome component | Both the 26S proteasome core and regulatory particles were implicated to be involved in plant defense against pathogen infections. | flg22 | (Adams and Spoel, 2018; DIELEN et al., 2010; Lee et al., 2006; Marino et al., 2012; Üstün et al., 2016; Yao et al., |
| AT3G27430 | PBB1 | 26S proteasome subunit beta type-7A |  |  | flg22 |  |
| AT3G26340 | PBE2 | 26S proteasome subunit beta type-5B |  |  | flg22 |  |
| AT5G58290 | RPT3 | 26S<br>proteasome<br>regulatory<br>AAA-ATPase<br>subunit |  |  | flg22 | 2012) |
| AT5G20000 | RPT6b | 26S<br>proteasome<br>regulatory<br>AAA-ATPase<br>subunit |  |  | flg22 |  |
| AT2G32730 | RPN2a | 26S<br>proteasome<br>regulatory<br>subunit |  |  | flg22 |  |
| AT1G75990 | RPN3b | 26S<br>proteasome<br>regulatory<br>subunit |  |  | flg22 |  |
| AT1G29150 | RPN6 | 26S<br>proteasome<br>regulatory<br>subunit |  |  | flg22 |  |
| AT3G11270 | RPN8b | 26S<br>proteasome<br>regulatory<br>subunit |  |  | flg22 |  |
| AT1G53750 | RPT1a | 26S<br>proteasome<br>regulatory<br>AAA-ATPase<br>subunit |  |  | Control,<br>flg22 |  |
| AT1G45000 | RPT4b | 26S<br>proteasome<br>regulatory<br>AAA-ATPase<br>subunit |  |  | Control,<br>flg22 |  |
| AT2G20580 | RPN1a | 26S proteasome regulatory subunit |  |  | Control, flg22 |  |
| AT1G20200 | RPN3a, EMB2719 | 26S proteasome regulatory subunit |  |  | Control, flg22 |  |
| AT4G24820 | RPN7 | 26S proteasome regulatory subunit |  |  | Control, flg22 |  |
| AT5G05780 | RPN8a | 26S proteasome regulatory subunit |  |  | Control, flg22 |  |
| AT5G23540 | RPN11 | 26S proteasome regulatory subunit |  |  | Control, flg22 |  |
| AT1G09100 | RPT5b | 26S proteasome regulatory AAA-ATPase subunit |  |  | Control |  |
| AT1G04810 | RPN2b | 26S proteasome regulatory subunit |  |  | Control |  |
| AT2G02560 | CAND1, ETA2 | Cullin-associated and neddylation dissociated 1 | Ub component | Regulators of SCF (CPR1) complexes | flg22 | (Huang et al., 2016) |
| AT3G20620 |  | F-box family protein-related |  | Not known. | Control, flg22 |  |
| AT3G23633 |  | F-box associated ubiquitylation effector family protein |  | Not known. | Control, flg22 |  |
| AT1G77710 | Ufm1, CCP2 | Ub-like, Ufm1 |  | Not known. | Control |  |

A common fate for ubiquitylated proteins is turnover by the 26S proteasome. This multisubunit particle consists of a 20S cylinder-shaped core protease (CP) that houses the peptidase activities, which is capped at one or both ends by the 19S regulatory particle (RP) that assist in substrate recognition, poly-Ub chain removal, unfolding, and finally translocation of the unfolded protein into to the CP for digestion (Marshall and Vierstra, 2018; Zhou and Zeng, 2017). The CP is assembled from four stacked heptameric rings of related α (PAA-PAG) and β (PBA-PBG) subunits arranged in an αββα configuration. By contrast, RP is created by a ring of six RP ATPases (RPT1-RPT6) together with a collection of non-ATPase subunits (RPN1-3 and RPN5-14). The RPT ring provides the unfoldase activities that prepare substrates for CP import, RPN10, RPN13, and possible RPN14/SEM1 serves as Ub receptors, and RPN11 is one of the several proteasome-bound DUBs that help release the Ub moieties prior to proteolysis. Unexpectedly, we identified PAC1 (α3), PBB1 (β2), and PBE2 (β5) within the CP, RPT3, and RPT6b within the RP ATPase ring, and RPN2b, RPN3a, RPN6, and RPN8a/b in the RP cap, as part of the flg22-induced ubiquitylome (Figure 2B; Tables 1 and 2). Additional RPN subunits were identified (RPT1a, RPT4b, RPT5b, RPN1a, RPN7, and RPN11) in the ubiquitylome from untreated samples as well. While this extensive modification of proteasomes is consistent with its ubiquitylation helping remove damaged particles via autophagy more generally (Marshall et al., 2015), and possibly as a target of during pathogen attack more specifically (Ustin et al., 2017), the preferential and rapid modification of some subunits after flg22 treatment also suggested a direct host role for this modification during the immune defense. As the selective modifications of several proteasome subunits occurred within 30 min of flg22 treatment and long before any impact on 26S proteasome synthesis, we propose that they reflect early defense signaling events.

### Flg22 induces UBC13 ubiquitylation and its role in plant immunity

We further connected the ubiquitylation target UBC13 (AT3G46460) to flg22 perception by studying its ubiquitylation profile and impact on pathogen defense. (It should be mentioned that UBC13 is unrelated to the UBC35 and UBC36 subfamily in *Arabidopsis*, which is unfortunately also named UBC13A (At1g78870) and UBC13B (At1g16890), respectively (Wen et al., 2008).) As shown in Figure 6A, we confirmed that UBC13 is ubiquitylated using a UBC13-HA variant co-expressed in Arabidopsis leaf protoplasts with FLAG-Ub. Only a monoubiquitylated form was evident in the immunoprecipitates from both untreated and flg22-treated samples; intriguingly, its abundance was slightly elevated upon flg22 exposure. Whether this addition was directed by autoubiquitylation or by a second factor (E3?) is not currently known.

**Figure 6.**
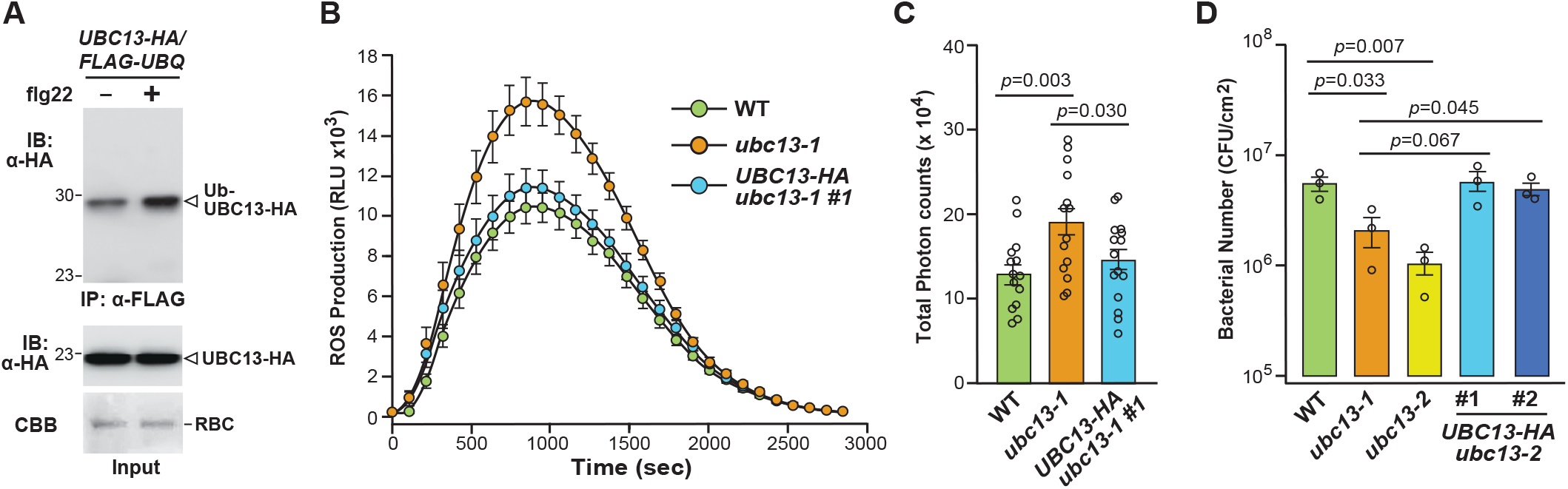
UBC13 is ubiquitylated and negatively regulates PTI responses. **A.** *In vivo* ubiquitylation of UBC13 upon flg22 treatment. Protoplasts were co-transfected with *FLAG-UBQ* and HA-tagged UBC13 (*UBC13-HA*) and incubated for 10 hr followed by treatment with 100 nM flg22 for 30 min. Protein extracts were immunoprecipitated with anti-FLAG beads (IP: α-FLAG) and the ubiquitylated proteins were immunoblotted with anti-HA antibodies (top). The input control was shown by an anti-HA immunoblot (bottom). **B, C.** UBC13 negatively regulates flg22-triggered ROS production. Leaf disks of four-week-old WT, *ubc13* (*CS389049*/*ubc13-1* & *CS389056*/*ubc13-2*), and plants harboring the *p35S::UBC13-HA* transgene in the *ubc13-2* mutant background were treated with 100 nM flg22. ROS production was monitored over time. The data are shown as mean ± s.e.m overlaid on dot plot (n =16) of RLU. The total photon count was shown in (**C**). **D.** *ubc13* mutants have elevated resistance against *Pst* DC3000. Leaf disks of four-week-old soil-grown WT, *ubc13-1*, *ubc13-2,* and *p35S::UBC13-HA ubc13-2* (Line 1 and 2) plants were hand-infiltrated with *Pst* DC3000 at OD_600_=5×10^−4^ CFU/ml, and bacterial growth was counted at 3 dpi. The data are shown as mean ± s.e.m overlaid on the dot plot (n = 3). The experiments were performed three times with similar results.

To examine whether UBC13 influences immune signaling, we examined two T-DNA insertion mutants *ubc13-1* (CS389049) and *ubc13-2*, (CS389056) available from the GABI-Kat collection (Kleinboelting et al., 2012). Both mutants were predicted to harbor a T-DNA sequence within the 6^th^ exon of *UBC13,* which was confirmed by genomic PCR with gene-specific and T-DNA border primers (Figure S1B, C). When compared to wild-type *Arabidopsis* Col-0 leaves, homozygous *ubc13-2* seedlings displayed enhanced ROS production upon flg22 treatment, based on the chemiluminescence assay, and were significantly more resistant to infection spread by the pathogen *Pst* DC3000 (Figure 5D). Confirmation that the *ubc13* mutations were responsible was provided using complementation studies that attempted to rescue the *ubc13-2* phenotypes with a transgene expressing UBC13-HA. Both ROS production and infectivity by the *Pst* DC3000 pathogen was returned to wild-type levels in homozygous *UBC13-HA ubc13-2* plants (Figure 6B-D). Together, our results imply that UBC13 negatively regulates plant immunity possibly by increasing the ubiquitylation state of UBC13.

### Flg22 induces polyubiquitylation of RPN8b to potentially impact plant immunity

Our discovery that RPN8 is rapidly ubiquitylated upon treating Arabidopsis with flg22 was particularly intriguing because of its role in RP assembly where it associates with the DUB RPN11 to form an RPN11-RPN8 hetero-dimeric subcomplex (Murata et al., 2009; Smalle and Vierstra, 2004). Two paralogs of RPN8 (RPN8a, and RPN8b) assemble into the 26S particle in *Arabidopsis*. While the RPNA8a isoform was more highly expressed based on mRNA levels and more commonly detected within the particle by MS (Book et al., 2010), only RPN8b was identified here in flg22-treated ubiquitylome, suggesting that its modification has a selective effect on 26S proteasome structure/function. As shown in Figure 7A, we confirmed the ubiquitylation of RPN8b by co-expressing an HA-tagged variant with FLAG-Ub in our *Arabidopsis* protoplast ubiquitylation system. Polyubiquitylated species of RPN8b-HA were evident in the immunoprecipitates from both untreated and flg22-treated samples, but their abundance was markedly enhanced upon flg22 treatment (Figure 7A).

**Figure 7.**
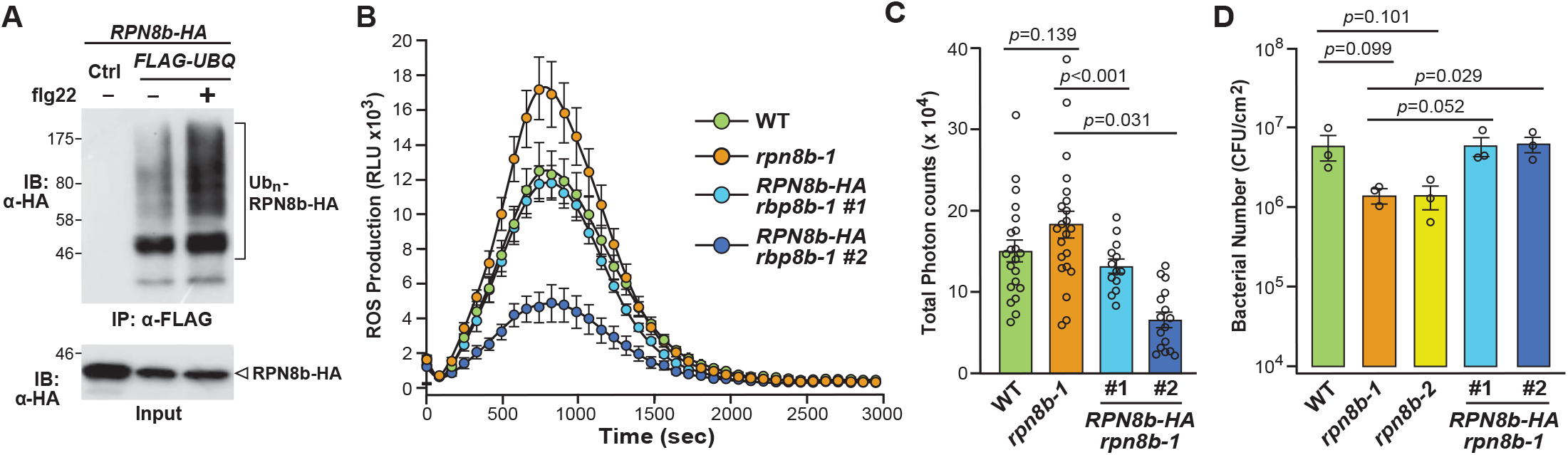
RPN8b is ubiquitylated and negatively regulates PTI responses. **A.** *in vivo* ubiquitylation of RPN8b upon flg22 treatment. Protoplasts were co-transfected with *FLAG-UBQ* and HA-tagged RPN8b (*RPN8b-HA*) or a control vector (Ctrl) and incubated for 10 hr followed by treatment with 100 nM flg22 for 30 min. Protein extracts from protoplasts were immunoprecipitated with anti-FLAG beads (IP: α-FLAG) and the ubiquitylated proteins were immunoblotted with anti-HA antibodies (top). The input control was shown by an anti-HA immunoblot (bottom). **B, C.** RPN8b negatively regulates flg22-triggered ROS production. Leaf disks of four-week-old WT, *rpn8b-1* (*SALK_128568*)*, rpn8b-2* (*SALK_023568*), and *rpn8b-1* plants harboring the *p35S::RPN8b-HA* transgene were treated with 100 nM flg22 or water as a control, and ROS production was monitored over time. The data are shown as mean ± s.e.m overlaid on the dot plot (n ≥ 13) of RLU. The total photon count was shown in (**C**). **D.** *rpn8b* mutants have elevated resistance against *Pst* DC3000. Leaf disks from four-week-old soil-grown WT, *rpn8b-1, rpn8b-2*, and *35S::RPN8b rpn8b-1* plants were hand-infiltrated with *Pst* DC3000 at OD_600_=5×10^−4^ CFU/ml, and bacterial growth was counted at 3 dpi. The data are shown as mean ± s.e.m overlaid on dot plot (n = 3). The experiments were performed three times with similar results.

To test whether RPN8b contributes to immune signaling, we analyzed two T-DNA insertion mutants available from the SALK insertion collection (*rpn8b-1*, *SALK_128568* and *rpn8b-2*, *SALK_023568*). The *rpn8b-1* allele was previously shown to harbor a T-DNA sequence within the 1^st^ intron (Palm et al., 2019), whereas we found that the *rpn8b-2* allele harbored a T-DNA sequence within the 2^nd^ intron, both of which were confirmed here by genomic PCR with gene-specific and T-DNA border primers (Figure S1D). When compared to wild-type *Arabidopsis* Col-0 leaves, homozygous *rpn8b-1* mutant leaves showed an increase in ROS production upon flg22 perception by the chemiluminescence assay, and enhanced disease resistance upon infiltrating leaves with *Pst* DC3000 (Figure 7B-D). Complementation studies then connected RPN8b to the phenotypes described here; transgenic plants expressing RPN8b-HA under the control of the CaMV *35S* promoter in the *rpn8b-1* background had their levels of flg22-induced ROS production and disease resistance restored to those in wild type (Figure 7B-D).

Collectively, the data implicated RPN8b as a negative regulator of plant immunity possibly through a process that enhances its ubiquitylation state. Whether the RPN8a isoform is similarly ubiquitylated and involved in disease resistance is not yet known. However, we note that suppression of ROS production in *rpn8b-1* seedlings by introducing *RPN8b-HA* was even more striking for one of the rescued lines, suggesting that overexpression of RPN8b might provide a dominant effect possibly by swamping out the function(s) of RPN8a (Figure 7B-D).

### Mapping ubiquitylation sites on Ub conjugates

One strategy to demonstrate the impact of ubiquitylation on protein function/stability is to identify the modified lysine within the target and then block this modification through lysine to arginine substitutions. Consequently, a catalog of such sites is valuable, which can be detected by MS identification of Ub footprints, *i.e.,* trypsin-protected lysines bearing an isopeptide linked di-glycine moiety (+114 kDa) (see Figure 8A). Directed queries of all of our MS datasets identified 150 ubiquitylation sites from 120 *Arabidopsis* proteins (Table 3), which adds to the expanding list generated by other reports (Aguilar-Hernandez et al., 2017; Kim et al., 2013; Maor et al., 2007; Saracco et al., 2009). Notable footprints within disease-related proteins identified here, some of which were previously unmapped, included K668 from CUL4, K301 from RPT6b, K502 and K507 from ARGONAUTE 7 (AGO7), K422 and K434 within the NLR proteins At1g63350 and At5g48770 respectively, K286 from UCH family protein At4g24320, and K736 from PHOSPHATIDYLINOSITOL-4-PHOSPHATE 5-KINASE (PIP5K2) (Table 3; Supplemental Table 2). Our datasets broadened the number of known Ub-attachment sites, especially within *Arabidopsis* proteins involved in flg22-triggered immune responses. Interestingly, noncanonical ubiquitylation sites detected in yeast or mammals, such as those involving serine, threonine, or cysteine residues (Iwai and Tokunaga, 2009; Okumoto et al., 2011; Shimizu et al., 2010) were not identified in here. Their absence is consistent with the previous ubiquitylome studies (Kim et al., 2013) and implies that these linkages are rarely, if at all, used by plants.

**Figure 8.**
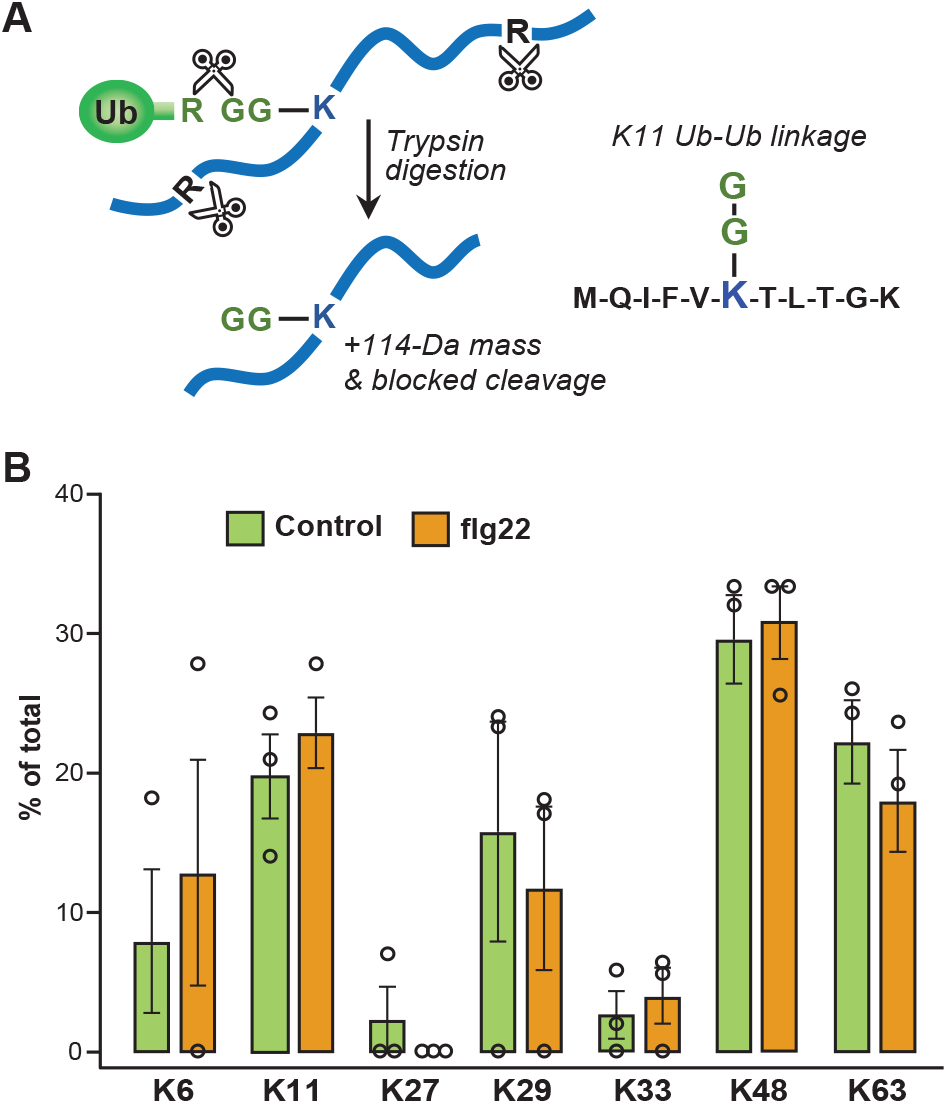
MS/MS mapping of Ub-Ub linkages. **A.** Strategy for detecting ubiquitylation sites by MS/MS. After trypsin digestion, a diglycine remnant of Ub covalently appended to a lysine residue of a conjugated protein is detected by an increased mass of 114 kDa for the lysine residue, along with protection from trypsin cleavage after that residue. Amino acids are denoted by single-letter code. **B.** The distribution of Ub-Ub linkages across the seven Ub lysines based on MS analysis from *Arabidopsis* transgenic plants carrying 6*HIS-UBQ* seedlings treated with or without 100 nM flg22. Percentage of Ub footprints at each site were obtained from PSM counts of diagnostic footprint peptides.

**Table 3.**
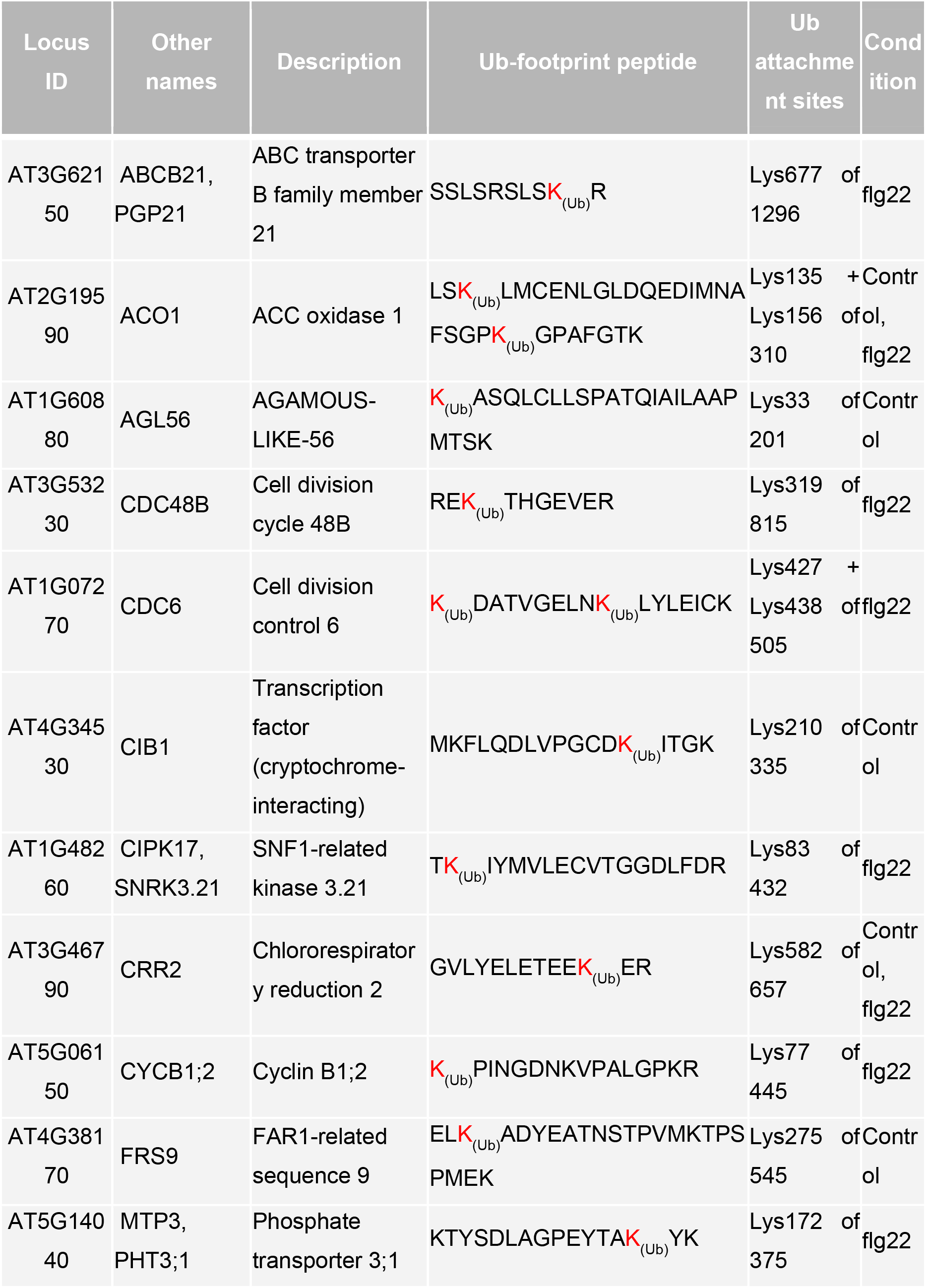
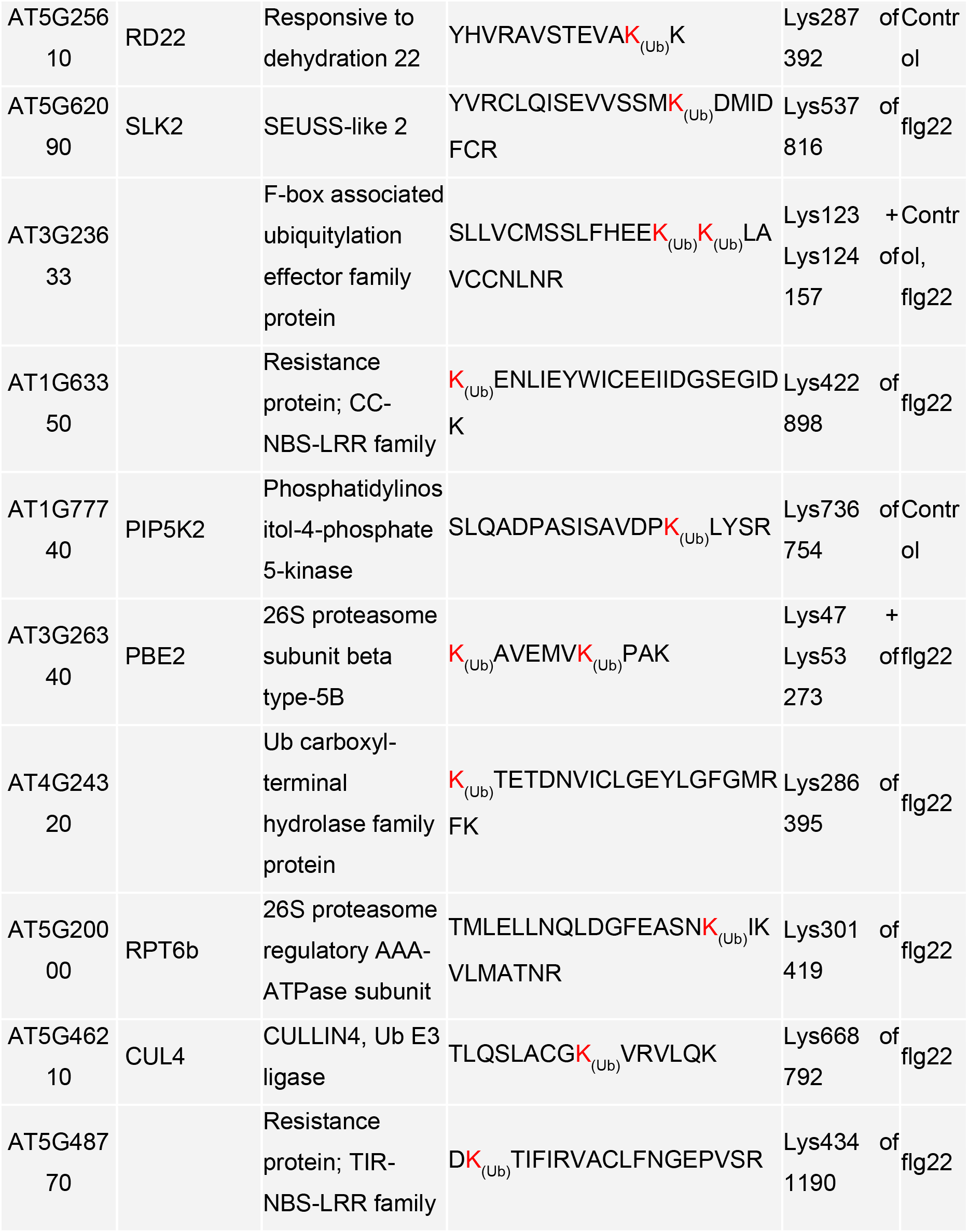
Examples of candidates with Ub-footprints identified from ubiquitylomes.

### Analysis of poly-Ub linkages

Owing to the widespread observations that poly-Ub chains of various linkages confer unique functions to Ub (Komander and Rape, 2012; Vierstra, 2009), we examined how flg22 treatment might affect the abundances of these polymers, using the diagnostic Ub footprint peptides for quantification based on peptide spectral matches (Peng et al., 2003). While footprints on each of the seven internal lysines (K^6^, K^11^, K^27^, K^29^, K^33^, K^48^, and K^63^) were identified (Figure 8B), none were seen using the N-terminal methionine, which would have signified linear Ub concatemers (Iwai and Tokunaga, 2009). K48-linked Ub-Ub connections were the most abundant and comprised ∼30% of the total detected linkages in both control and flg22-treated samples (Figure 8C). The second most abundant linkage involved K63 and was followed in abundance by those involving K11, K29, K6, K33, and K27 (Figure 8B). Interestingly, flg22 exposure altered the relative abundance of poly-Ub linkages; the percentages of K6, K11, K33, and K48-linkages increased, while the percentages of K27, K29, and K63-linkages decreased (Figure 8B). The biological relevance of such changes awaits clarification.

## DISCUSSION

Protein ubiquitylation is involved in nearly all aspects of eukaryotic biology, leading to a multitude of distinct signals within a variety of cellular and physiological contexts. In line with prior studies with yeast and mammals (Bennett et al., 2007; Ordureau et al., 2020), we show here and elsewhere (Aguilar-Hernandez et al., 2017) that dynamic and multifaceted changes in the plant ubiquitylome also occur. By employing a genetically encoded *hexa*-*6HIS-UBQ* variant coupled with an improved two-step enrichment protocol for ubiquitylated proteins, followed by deep LC-MS/MS analysis, we found profound and complex protein ubiquitylation patterns upon immune elicitation in Arabidopsis.

We propose that the use of two affinity-purification steps, with the second under strongly denaturing conditions, are essential to generate reliable catalogs, which resulted here in the detection of 961 possible substrates along with 150 ubiquitylation sites mapped onto 120 individual proteins. Among the 10 candidate proteins that we subsequently tested by *in vivo* conjugation assays, all were confirmed as ubiquitylated targets *in planta*. While it is possible that the ubiquitylated species seen in protoplasts were artifacts derived from ectopic expression, the fact that different types of linkages were seen (mono-versus polyubiquitylation) and that both increased and decreased responses were seen soon after flg22 elicitation argue otherwise.

We also identified Ub-Ub linkages involving all seven lysine residues based on the discovery of diagnostic footprint peptides, with those involving K48, K63, and K11 being the most abundant. Whereas K48 polyubiquitylation has been linked to 26S proteasome-mediated turnover, and K63 ubiquitylation has been connected to the endocytic internalization of plasma membrane proteins (Romero-Barrios and Vert, 2018), the functions of the other poly-Ub linkages remain largely unknown in *Arabidopsis* even though the total abundance of those linkages exceeds 40% of the total. In addition, we saw an increased number of ubiquitylated proteins (391 of control vs. 570 of flg22-treated) upon immune elicitation. Although the datasets were not derived from quantitative proteomic approaches, a nearly 50% increase in the number of ubiquitylated proteins seen soon after immune elicitation, suggests a strong activation of the machinery that supports Ub transfer.

Notably, our flg22-induced datasets were obtained using seedlings treated for only 30 minutes, thus implicating ubiquitylation as an early step in immune signaling. Surprisingly, several resistance proteins known to be ubiquitylation such as FLS2, BRI1, and BIK1 were not identified here (Lu et al., 2011; Ma et al., 2020; Zhou et al., 2018), which might be explained by the nature of the tissues examined or the extraction method used, which possibly preferred cytoplasmic versus plasma membrane/organelle-associated proteins. Furthermore, we should also emphasize the TUBE affinity approach likely binds polyubiquitylated proteins better than monoubiquitylated proteins and thus reduced the latter in our catalogs. One strategy to avoid this complication is to enrich for Ub footprints directly from trypsin-digested cell lysates using an anti- K-ԑ-GG antibodies specific for the GG-remnant on ubiquitylated lysines (Udeshi et al., 2013). In fact, recent studies with such antibodies in rice discovered 1,376 possible ubiquitylated proteins and revealed strong changes in ubiquitylation after a three-hr exposure to flg22 or fungal chitin (Chen et al., 2018). The only complications were that designation of a protein as an Ub target was often based on a single Ub footprint peptide and the reliability of the anti-K-ԑ-GG antibodies. Clearly, further advancements in enrichment strategies and MS instrumentation should further expand the specificity, stringency, and depth of ubiquitylome during diverse physiological conditions.

Within the flg22-induced ubiquitylomes identified here, we confirmed several Ub targets by *in vivo* ubiquitylation assays in protoplasts. Surprisingly, we found distinct ubiquitylation patterns and dynamic temporal responses to flg22 treatment. For example, MKK5 and MPK3 were strongly monoubiquitylated while SOBIR1 and EDS1 were mainly polyubiquitylated. Ubiquitylation of RPN8b and MPK3 was increased by flg22, with MPK3 modification showing a transient effect. In contrast, ubiquitylation of EDS1 in the protoplasts was reduced in response to flg22 treatment. One of the more interesting substrates identified here was receptor protein kinase RKL1, which was polyubiquitylated, and found genetically to negatively regulate flg22-triggered ROS and immunity against bacterial *Pst* DC3000 infection. Intriguingly, RLK902, a close homolog of RKL1, was previously shown to positively regulate plant disease resistance (Zhao et al., 2019), suggesting that RKL1 and RLK902 counterbalance each other in immune signaling.

Ubiquitylation often intertwines with phosphorylation in regulating protein function (Hu and Sun, 2016; Reyes et al., 2011; Skubacz et al., 2016). A well-known example in plant disease resistance is BIK1, which is phosphorylated and subsequently monoubiquitylated upon flg22 perception (Ma et al., 2020). Here, we found that the phosphorylation (Asai et al., 2002) target MPK3 is also ubiquitylated dynamically, and like phosphorylation, this modification was enhanced ∼10 min after flg22 treatment, and then was reduced to basal levels after 30 min. The cognate E3 ligase for MPK3 is currently unknown; given that MPK3 phosphorylates the E3 ligase PUB22 and regulates its turnover (Furlan et al., 2017), it is possible that PUB22 directs MPK3 ubiquitylation thus creating an internal regulatory loop. As MPK3 remained unchanged despite the dynamics of its ubiquitylation after flg22 treatment, its ubiquitylation might not commit MPK3 to turnover but confer a reversible, non-proteolytic function. In line with some ubiquitylation events being reversible, we note the detection of several DUBs in our flg22-treated ubiquitylome datasets.

Pathway analysis of our ubiquitylome datasets upon MAMP perception revealed robust changes in the ubiquitylation status for numerous components involved in the UPS, including E1, E2, E3, and DUB family members. The *in vivo* ubiquitylation assays further confirmed that the E2 UBC13 in monoubiquitylated *in vivo* with the abundance of this modification increased upon flg22 treatment. While *ubc13* mutants morphologically resembled WT plants, they displayed an enhanced flg22-induced ROS burst and increased resistance against bacterial infection. Several E3s have also been implicated in immunity, including CPR1 (Cheng et al., 2011; Gou et al., 2012; Gou et al., 2009), PUB12/13 (Liao et al., 2017; Lu et al., 2011), PUB25/26 (Wang et al., 2018), PUB22/23/24 (Stegmann et al., 2012), and RHA3A/B (Ma et al., 2020). Although it is generally accepted that E3s, not E2s, are the key determinants in substrate selection, it is possible that UBC13 preferentially works with those E3s involved in immunity.

Proteasome components, including both the core and regulatory particles, are enriched in our ubiquitylome analysis, which is consistent with the observation in yeast (Peng et al., 2003), and the known role of ubiquitylation in directing proteasome turnover through autophagy (Marshall et al., 2015). Strikingly, several of the ubiquitylated subunits were only found in the flg22-treated ubiquitylome, implying an active regulation of the proteasome machinery upon flg22 perception. Previous studies showed that several regulatory particle subunits, such as RPN1a, RPN8a, and RPT2a, but not others, play a positive role in immunity (Yao et al., 2012). Unexpectedly, we found RPN8b negatively regulates the flg22-induced ROS burst and disease resistance against *Pst* DC3000 infection. As Arabidopsis also encodes a second isoform of RPN8 (RPN8a) that appears more dominant in providing this subunit (Book et al., 2010), one intriguing possibility is that the RPN8b isoform becomes assembled to a unique subset of proteasomes with functions more directed toward immune perception. The immunoproteasome assembled with unique isoforms of the CP subunits β1, β2, and β5 could serve as an example of such non-redundancy (Finley, 2009). Clearly, further comparisons of mutants impacting both RPN8a and RPN8b with respect to pathogen resistance, along with assays testing the ubiquitylation status of each is now necessary to explore this notion.

## Acknowledgements

We thank the *Arabidopsis* Biological Resource Center (ABRC) for T-DNA insertion mutant seeds, and members of the laboratories of L.S., P.H. and R.D.V. for discussions and comments of the experiments. The work was supported by grants from the National Institutes of Health (NIH) (R01-GM092893) and National Science Foundation (NSF) (MCB-1906060) to P.H., NIH (R01-GM097247) and the Robert A. Welch Foundation (A-1795) grant to L.S., and a grant from the NIH (R01-GM124452) to R.D.V. C. Z. and Y.H. were partially supported by the China Scholarship Council (CSC).

## Author Contributions

R.D.V., P.H., and L.S. conceived the project, designed experiments, and analyzed data. X.M., C.Z., D.K., and Y.H. performed experiments and analyzed data. X.M., C.Z., R.D.V., and L.S. wrote the manuscript with inputs from all co-authors.

## MATERIALS AND METHODS

### Plant materials and growth conditions

*Arabidopsis thaliana 6HIS-UBQ* transgenic plants in the Col-0 background were described previously (Saracco et al., 2009). The *rkl1-1* (*SALK_099094*), *ubc13-1* (*CS389049*), *ubc13-2* (*CS389056*), *rpn8b-1* (*SALK_128568*), *rpn8b-2* (*SALK_023568*) T-DNA insertion lines were obtained from ABRC. *p35S::UBC13-HA* transgenic plants in the *ubc13-2* background and *p35S::RPB8b-HA* transgenic plants in the *rpn8b-1* background were generated in this study (see below for details). All *Arabidopsis* plants were grown in soil (Metro Mix 366, Sunshine LP5 or Sunshine LC1) in a growth chamber at 20-23 °C, 50% relative humidity and 75 μE m^− 2^s^−1^ light with a 12-hr light/12-hr dark photoperiod for four weeks before pathogen infection assay, protoplast isolation, and ROS assays.

### Plasmid construction and generation of transgenic plants

*MPK3-HA*, *MKK5-MYC*, and *FLAG-UBQ* in a plant gene expression vector *pHBT* used for protoplast assays were described previously (Asai et al., 2002; Lu et al., 2011). SOBIR1, RKL1, UBC13, RPN8b, and EDS1 tagged with HA in the *pHBT* vector were generated as following: the gene was PCR amplified from total Col-0 cDNA with primers containing *Bam*HI (for *SOBIR1*, *RKL1*, *UBC13*, and *RPN8b*) or *Nco*I (for *EDS1*) at the 5’ end and *Stu*I at the 3’ end, followed by *Bam*HI/*Nco*I and *Stu*I digestion and ligation into the *pHBT* vector with a sequence encoding a HA tag at the 3’ end. DNA fragments cloned into the *pHBT* vectors were confirmed as correct by Sanger sequencing. The *UBC13-HA* and *RPN8b-HA* transgenes were further shuttled into a *pCB302* vector by insertion into *Bam*HI and *Stu*I digestion sites. The *A. tumefaciens*-mediated floral dip was used to transform the above binary vectors into *ubc13-2* or *rpn8b-1* plants. Transgenic plants were selected by glufosinate-ammonium (Basta, 50 μg/ml) resistance. Multiple transgenic lines were analyzed by immunoblotting for protein expression. Two lines with a 3:1 segregation ratio for Basta resistance in the T3 generation were selected to obtain homozygous seeds for further studies.

### Affinity purification of ubiquitylated proteins

Affinity purification of ubiquitylated proteins was described (Kim et al., 2013) with modifications. Seeds of *Arabidopsis* Col-0 and *6HIS-UBQ* transgenic plants were vernalized at 4 °C for three days and surface-sterilized with 75% ethanol for 10 min and 5% bleach for 10 min, and plated in 1/2-strength Murashige and Skoop (MS) medium containing 1% sucrose and 0.5% agar (100 seedlings per plate) at 23 °C, 75 μE m^−2^s^−1^ light with a 12-hr light/12-hr dark photoperiod for 14 days. Approximately 5000 seedlings (50 plates) were used for each biological replicate which generated ∼16-21 gram fresh weight of tissue. Seedlings were gently transferred into a Petri dish plate and incubated overnight in water. After removing the water by vacuum, seedlings were treated for 30 min with 100 nM flg22 or water as a control.

Seedlings were gently blot-dried on Kimwipes, frozen to liquid nitrogen temperatures, and homogenized in 0.5 g/mL of extraction buffer (EB; 50 mM Tris-HCl, pH 7.2, and 200 mM NaCl), containing 0.25% Triton X-100, 13 protease inhibitor cocktail (Roche, 1 tablet per 10 ml of EB), 2 mM phenylmethanesulfonyl fluoride, 10 mM 2-chloroacetamide, 10 mM sodium metasulfite, and 1 mM N-ethylmaleimide (freshly added). The homogenate was filtered through two layers of Miracloth and the supernatant was mixed with TUBEs beads as the ratio of 40 g of tissue to 1 mL of beads. After incubated for 6 hr at 4°C, the beads were collected in a 2 × 1-0cm chromatography column and washed three times with EB and three times with EB plus 1, 2, 4 M NaCl, and eluted in 10 mL of 7 M guanidine-HCl, 100 mM NaH_2_PO_4_, and 10 mM Tris-HCl (pH 8.0, 24 °C). The eluted conjugates were incubated with 500 µl of Ni-NTA beads for 12 hr at 4°C in the same buffer with the addition of 20 mM imidazole and 10 mM 2-chloroacetamide. The beads were collected and washed once in 6 M guanidine-HCl, 0.1% SDS, 100 mM NaH_2_PO_4_, and 10 mM Tris-HCl (pH 8.0), and once in urea buffer (UB; 8 M urea, 100 mM NaH_2_PO_4_, and 10 mM Tris-HCl (pH 8.0)) also containing 0.1% Triton X-100, twice in UB plus 20 mM imidazole, and three times in UB alone, and eluted by 1 ml of UB supplemented with 400 mM imidazole. The two-step purified conjugates were filtered by the Ultracel-10K filter (Millipore) and separated by SDS-PAGE followed by staining for protein with silver, or immunoblotting with anti-Ub (Sigma-Aldrich) or anti-His (Novagen) antibodies to detect Ub conjugates.

### Mass spectrometry

Purified proteins (100 μl) or control samples were reduced with 10 mM DTT at room temperature for 1 hr, and alkylated in the dark in the presence of 30 mM 2-chloroacetamide at room temperature for a further 1 hr. Excess alkylating agent was quenched with 20 μL of 200 mM DTT for 10 min. Samples, ten-fold diluted with 25 mM ammonium bicarbonate, were digested for 12 hr at 37 °C with 2 mg of sequencing-grade trypsin (Promega), followed by a second incubation for 6 hr with an additional 2 mg of trypsin. Digestion products were desalted using a C18 solid-phase extraction pipette tip (SPEC PT C18, Varian), vacuum dried, and reconstituted in 10 μL of 0.1% formic acid and 5% acetonitrile in water for MS analysis.

Samples were analyzed by a nanoflow liquid chromatography system (nanoAcquity; Waters) combined with an electrospray ionization FT/ion-trap mass spectrophotometer (LTQ Orbitrap Velos; Thermo Fisher Scientific). The LC system employed a 100 × 365 -μm fused silica microcapillary column packed with 15 cm of 3-μm-diameter, 100-Å pore size, C18 beads (Magic C18; Bruker), with the emitter tip pulled to 2 mm using a laser puller (Sutter Instruments). Peptides were loaded onto the column for 30 min at a flow rate of 500 nL/ min and eluted over 120 min at a 200 nL/min flow rate with a gradient of 2% to 30% acetonitrile in 0.1% formic acid. MS spectra were acquired in the FT orbitrap between 300 and 1500 mass-to-charge ratios (m/z) at a resolution of 60 000, followed by 10 MS/MS HCD scans of the 10 highest intensity parent ions at 42% relative collision energy and 7500 resolution, with a mass range starting at 100 mass-to-charge ratios. Dynamic exclusion was enabled with a repeat value of two for 30 sec and an exclusion window for 120 sec.

### MS data analysis

The MS/MS data were searched using SEQUEST version 1.2 (ThermoFisher Scientific) against the *A. thaliana* ecotype Col-0 protein database (IPI database, version 3.85, containing 39,677 entries available within the *Arabidopsis* Information Resource [TAIR], http://www.arabidopsis.org) databse. Masses for both precursor and fragment ions were treated as monoisotopic. Met oxidation (+15.994915 Da), Cys carbamidomethylation (+57.021464 Da), the di-Gly Ub footprint indicating Ub addition to a lysine residue (+114.043 Da) were set as variable modifications. The database search allowed for up to two missed trypsin cleavages, and the ion mass tolerances were set to 10 ppm for precursor and 0.1 Da for HCD fragments. The data were filtered using a 1% FDR, and a minimum of two peptide matches was required or at least one peptide if it included a GGK Ub footprint.

GO analysis was evaluated by DAVID Functional Annotation Tool (Huang da et al., 2009) using GO annotations in the TAIR GO database (http://www.arabidopsis.org/). Fold enrichments were calculated based on the frequency of proteins annotated to the term compared with their frequency in the proteome. The *p*-value combined with the FDR correction was used as criteria of significant enrichment for GO catalogs, whereas a *p*-value < 0.05 were considered to be enriched for GO terms. The GO annotation was classified based on the “biological processes,” “molecular functions,” and “cellular components” categories and visualized as a bubble plot by R Project 4.0.0 (https://www.r-project.org/) (Bonnot et al., 2019). Protein interaction networks were created by the STRING database version 11.0 (Szklarczyk et al., 2015), and visualized by Cytoscape version 3.8.0 (Shannon et al., 2003).

### Detection of ROS production

Leaves from 4-5-week-old, soil-grown *Arabidopsis* plants were punched into 5-mm diameter discs. The discs were incubated in 100 μl of water with gentle shaking overnight, which was then replaced with 100 μl of reaction solution containing 50 μM luminol and 10 μg/ml horseradish peroxidase (Sigma-Aldrich) supplemented with or without 100 nM flg22. Luminescence was measured with a luminometer (GloMax®-Multi Detection System, Promega) with a setting of 1 min as the interval for 40 min. Detected values for ROS production were indicated as means of Relative Light Units (RLU).

### Pathogen infection assays

*Pseudomonas syringae* pv. *tomato* (*Pst*) DC3000 was cultured for overnight at 28 °C in King’s B medium supplemented with rifamycin (50 μg/ml). Cells were collected by centrifugation at 3,500 rpm, washed, and re-suspended to the density of 5 × 10^5^ cfu/ml in 10 mM MgCl_2_. Leaves from four-week-old *Arabidopsis* plants were hand-inoculated with bacterial suspension using a needleless syringe. To measure *in planta* bacterial growth, three to four samples, each containing two 6-mm diameter leaf discs, were homogenized in 100 μl of water, and plated in serial dilutions on medium containing 1% tryptone, 1% sucrose, 0.1% glutamic acid, 1.8% agar, and 25 μg/ml rifamycin. Plates were incubated at 28 °C and bacterial colony forming units (cfu) were counted after 2 days.

### *In vivo* ubiquitylation assays

Protoplast isolation and transient expression assay were as described previously (Zhou et al., 2014). Protoplasts were freshly isolated from leaves of four-week-old Col-0 plants and transfected at 2 × 10^5^ cell/ml (500 μL) with various combinations of plasmids harboring genes encoding FLAG-UBQ (25 μL at 2 µg DNA/µL), and the HA or MYC-tagged proteins of interested (25 μL at 2 µg DNA /µL). The protoplasts were incubated for overnight at room temperature followed by a 2-hr treatment with DMSO alone or 2 μM MG132 dissolved in DMSO, and then a 2-hr exposure to 100 nM flg22. After homogenization in 300 μL of Immunoprecipitation (IP) buffer (150 mM NaCl, 50 mM Tris-HCl (pH 7.5), 5 mM Na_2_EDTA, 1% Triton X-100, 2 mM Na_3_VO_4_, 2 mM NaF, 1 mM DTT, and 1:100 diluted protease inhibitor cocktail (Sigma-Aldrich), the FLAG-tagged proteins were immunoprecipitated by incubation of the extracts with 5 μL of anti-FLAG antibody agarose beads (Sigma-Aldrich) for 1 hr at 4 °C with gentle shaking. The anti-FLAG antibody beads were collected and washed three times with IP buffer (without cocktail) and once with 50 mM Tris-HCl (pH 7.5), and then incubated in 30 μL of hot SDS-PAGE sample buffer for 5 min. Twenty μL of the sample before adding the beads was used as the input control. The samples were separated by SDS-PAGE followed by electrophoretic transfer onto Immun-Blotpolyvinylidene difluoride (PVDF) membranes (Bio-Rad), and immunoblotting with appropriate antibodies. Antibodies against plant Ub (van Nocker et al., 1996) and 6His (Kim et al., 2013) were as described. Anti-FLAG, anti HA, and, anti-myc antibodies were purchased from Sigma-Aldrich (A8592), Roche (12013819001), and Santa Cruz (sc-40), respectively.

### Statistical analyses

Data for quantification analyses are presented as mean ± standard error of the mean (s.e.m.). The statistical analyses were performed by Student’s *t*-test or one-way analysis of variance (ANOVA) test (* *p*-value < 0.05, ** *p*-value < 0.01, *** *p*-value < 0.001). The number of replicates is shown in the figure legends.

## References

Adams, E.H.G., and Spoel, S.H. (2018). The ubiquitin-proteasome system as a transcriptional regulator of plant immunity. J Exp Bot 69, 4529–4537.

Aguilar-Hernandez, V., Kim, D.Y., Stankey, R.J., Scalf, M., Smith, L.M., and Vierstra, R.D. (2017). Mass Spectrometric Analyses Reveal a Central Role for Ubiquitylation in Remodeling the Arabidopsis Proteome during Photomorphogenesis. Mol Plant 10, 846–865.

Albert, I., Böhm, H., Albert, M., Feiler, C.E., Imkampe, J., Wallmeroth, N., Brancato, C., Raaymakers, T.M., Oome, S., and Zhang, H. (2015). An RLP23–SOBIR1–BAK1 complex mediates NLP-triggered immunity. Nature Plants 1, 1–9.

Alonso, J.M., Stepanova, A.N., Leisse, T.J., Kim, C.J., Chen, H., Shinn, P., Stevenson, D.K., Zimmerman, J., Barajas, P., Cheuk, R., et al. (2003). Genome-wide insertional mutagenesis of Arabidopsis thaliana. Science 301, 653–657.

Asai, T., Tena, G., Plotnikova, J., Willmann, M.R., Chiu, W.L., Gomez-Gomez, L., Boller, T., Ausubel, F.M., and Sheen, J. (2002). MAP kinase signalling cascade in Arabidopsis innate immunity. Nature 415, 977–983.

Bakker, E.G., Toomajian, C., Kreitman, M., and Bergelson, J. (2006). A genome-wide survey of R gene polymorphisms in Arabidopsis. Plant Cell 18, 1803–1818.

Bellati, J., Champeyroux, C., Hem, S., Rofidal, V., Krouk, G., Maurel, C., and Santoni, V. (2016). Novel Aquaporin Regulatory Mechanisms Revealed by Interactomics. Mol Cell Proteomics 15, 3473–3487.

Bennett, E.J., Shaler, T.A., Woodman, B., Ryu, K.Y., Zaitseva, T.S., Becker, C.H., Bates, G.P., Schulman, H., and Kopito, R.R. (2007). Global changes to the ubiquitin system in Huntington’s disease. Nature 448, 704–708.

Bethke, G., Pecher, P., Eschen-Lippold, L., Tsuda, K., Katagiri, F., Glazebrook, J., Scheel, D., and Lee, J. (2012). Activation of the Arabidopsis thaliana mitogen-activated protein kinase MPK11 by the flagellin-derived elicitor peptide, flg22. Mol Plant Microbe Interact 25, 471–480.

Böhm, H., Albert, I., Fan, L., Reinhard, A., and Nürnberger, T.J.C.o.i.p.b. (2014). Immune receptor complexes at the plant cell surface. 20, 47–54.

Bonnot, T., Gillard, M., and Nagel, D. (2019). A Simple Protocol for Informative Visualization of Enriched Gene Ontology Terms. Bio-Protocol 9.

Book, A.J., Gladman, N.P., Lee, S.S., Scalf, M., Smith, L.M., and Vierstra, R.D. (2010). Affinity purification of the Arabidopsis 26 S proteasome reveals a diverse array of plant proteolytic complexes. The Journal of biological chemistry 285, 25554–25569.

Chen, X.L., Xie, X., Wu, L., Liu, C., Zeng, L., Zhou, X., Luo, F., Wang, G.L., and Liu, W. (2018). Proteomic Analysis of Ubiquitinated Proteins in Rice (Oryza sativa) After Treatment With Pathogen-Associated Molecular Pattern (PAMP) Elicitors. Front Plant Sci 9, 1064.

Chen, Y., Li, F., Tian, L., Huang, M., Deng, R., Li, X., Chen, W., Wu, P., Li, M., and Jiang, H. (2017). The phenylalanine ammonia lyase gene LJPAL1 is involved in plant defense responses to pathogens and plays diverse roles in Lotus japonicus-rhizobium symbioses. Molecular Plant-Microbe Interactions 30, 739–753.

Cheng, Y.T., and Li, X. (2012). Ubiquitination in NB-LRR-mediated immunity. Curr Opin Plant Biol 15, 392–399.

Cheng, Y.T., Li, Y.Z., Huang, S.A., Huang, Y., Dong, X.N., Zhang, Y.L., and Li, X. (2011). Stability of plant immune-receptor resistance proteins is controlled by SKP1-CULLIN1-F-box (SCF)-mediated protein degradation. P Natl Acad Sci USA 108, 14694–14699.

Cheng, Z., Li, J.F., Niu, Y., Zhang, X.C., Woody, O.Z., Xiong, Y., Djonovic, S., Millet, Y., Bush, J., McConkey, B.J., et al. (2015). Pathogen-secreted proteases activate a novel plant immune pathway. Nature 521, 213–216.

Clay, N.K., Adio, A.M., Denoux, C., Jander, G., and Ausubel, F.M. (2009). Glucosinolate metabolites required for an Arabidopsis innate immune response. Science 323, 95–101.

Collins, N.C., Thordal-Christensen, H., Lipka, V., Bau, S., Kombrink, E., Qiu, J.L., Huckelhoven, R., Stein, M., Freialdenhoven, A., Somerville, S.C., et al. (2003). SNARE-protein-mediated disease resistance at the plant cell wall. Nature 425, 973–977.

Couto, D., and Zipfel, C. (2016). Regulation of pattern recognition receptor signalling in plants. Nat Rev Immunol 16, 537–552.

Cui, H., Tsuda, K., and Parker, J.E. (2015). Effector-triggered immunity: from pathogen perception to robust defense. Annual review of plant biology 66, 487–511.

Dielen, A.S., Badaoui, S., Candresse, T., and German-Retana, S. (2010). The ubiquitin/26S proteasome system in plant–pathogen interactions: a never-ending hide-and-seek game. Molecular plant pathology 11, 293–308.

Eschen-Lippold, L., Bethke, G., Palm-Forster, M.A., Pecher, P., Bauer, N., Glazebrook, J., Scheel, D., and Lee, J. (2012). MPK11—a fourth elicitor-responsive mitogen-activated protein kinase in Arabidopsis thaliana. Plant signaling & behavior 7, 1203–1205.

Ewan, R., Pangestuti, R., Thornber, S., Craig, A., Carr, C., O’Donnell, L., Zhang, C., and Sadanandom, A. (2011). Deubiquitinating enzymes AtUBP12 and AtUBP13 and their tobacco homologue NtUBP12 are negative regulators of plant immunity. New Phytol 191, 92–106.

Finley, D. (2009). Recognition and processing of ubiquitin-protein conjugates by the proteasome. Annual review of biochemistry 78, 477–513.

Fuchs, R., Kopischke, M., Klapprodt, C., Hause, G., Meyer, A.J., Schwarzlander, M., Fricker, M.D., and Lipka, V. (2016). Immobilized Subpopulations of Leaf Epidermal Mitochondria Mediate PENETRATION2-Dependent Pathogen Entry Control in Arabidopsis. Plant Cell 28, 130–145.

Furlan, G., Nakagami, H., Eschen-Lippold, L., Jiang, X., Majovsky, P., Kowarschik, K., Hoehenwarter, W., Lee, J., and Trujillo, M. (2017). Changes in PUB22 Ubiquitination Modes Triggered by MITOGEN-ACTIVATED PROTEIN KINASE3 Dampen the Immune Response. The Plant cell 29, 726–745.

Furniss, J.J., Grey, H., Wang, Z., Nomoto, M., Jackson, L., Tada, Y., and Spoel, S.H. (2018a). Proteasome-associated HECT-type ubiquitin ligase activity is required for plant immunity. PLoS Pathog 14, e1007447.

Furniss, J.J., Grey, H., Wang, Z., Nomoto, M., Jackson, L., Tada, Y., and Spoel, S.H. (2018b). Proteasome-associated HECT-type ubiquitin ligase activity is required for plant immunity. PLoS pathogens 14, e1007447.

Furniss, J.J., Grey, H., Wang, Z., Nomoto, M., Jackson, L., Tada, Y., and Spoel, S.H.J.P.p. (2018c). Proteasome-associated HECT-type ubiquitin ligase activity is required for plant immunity. 14, e1007447.

Goritschnig, S., Zhang, Y., and Li, X. (2007). The ubiquitin pathway is required for innate immunity in Arabidopsis. The Plant Journal 49, 540–551.

Gou, M., Shi, Z., Zhu, Y., Bao, Z., Wang, G., and Hua, J. (2012). The F-box protein CPR1/CPR30 negatively regulates R protein SNC1 accumulation. Plant J 69, 411–420.

Gou, M., Su, N., Zheng, J., Huai, J., Wu, G., Zhao, J., He, J., Tang, D., Yang, S., and Wang, G.J.T.P.J. (2009). An F-box gene, CPR30, functions as a negative regulator of the defense response in Arabidopsis. 60, 757–770.

Gu, Y., and Innes, R.W. (2011). The KEEP ON GOING protein of Arabidopsis recruits the ENHANCED DISEASE RESISTANCE1 protein to trans-Golgi network/early endosome vesicles. Plant Physiol 155, 1827–1838.

Gu, Y., Zavaliev, R., and Dong, X.J.M.p. (2017). Membrane trafficking in plant immunity. 10, 1026–1034.

Guerra, D.D., and Callis, J. (2012). Ubiquitin on the move: the ubiquitin modification system plays diverse roles in the regulation of endoplasmic reticulum-and plasma membrane-localized proteins. Plant Physiol 160, 56–64.

Heidrich, K., Wirthmueller, L., Tasset, C., Pouzet, C., Deslandes, L., and Parker, J.E. (2011). Arabidopsis EDS1 connects pathogen effector recognition to cell compartment–specific immune responses. Science 334, 1401–1404.

Hjerpe, R., Aillet, F., Lopitz-Otsoa, F., Lang, V., England, P., and Rodriguez, M.S. (2009). Efficient protection and isolation of ubiquitylated proteins using tandem ubiquitin-binding entities. EMBO reports 10, 1250–1258.

Hou, S., Wang, X., Chen, D., Yang, X., Wang, M., Turra, D., Di Pietro, A., and Zhang, W. (2014). The secreted peptide PIP1 amplifies immunity through receptor-like kinase 7. PLoS pathogens 10, e1004331.

Hou, X., and Gao, Y. (2017). Investigation on the Interaction of Pseudomonas syringae Effector AvrPto with AtRabE1d GTPase. Protein and peptide letters 24, 661–667.

Hu, H.B., and Sun, S.C. (2016). Ubiquitin signaling in immune responses. Cell research 26, 457–483.

Huang da, W., Sherman, B.T., and Lempicki, R.A. (2009). Systematic and integrative analysis of large gene lists using DAVID bioinformatics resources. Nature protocols 4, 44–57.

Huang, J., Gu, M., Lai, Z., Fan, B., Shi, K., Zhou, Y.H., Yu, J.Q., and Chen, Z. (2010). Functional analysis of the Arabidopsis PAL gene family in plant growth, development, and response to environmental stress. Plant Physiol 153, 1526–1538.

Huang, S., Chen, X., Zhong, X., Li, M., Ao, K., Huang, J., and Li, X. (2016). Plant TRAF Proteins Regulate NLR Immune Receptor Turnover. Cell Host Microbe 20, 271.

Isono, E., and Nagel, M.K. (2014). Deubiquitylating enzymes and their emerging role in plant biology. Front Plant Sci 5, 56.

Iwai, K., and Tokunaga, F. (2009). Linear polyubiquitination: a new regulator of NF-kappaB activation. EMBO reports 10, 706–713.

Jacobs, A.K., Lipka, V., Burton, R.A., Panstruga, R., Strizhov, N., Schulze-Lefert, P., and Fincher, G.B. (2003). An Arabidopsis callose synthase, GSL5, is required for wound and papillary callose formation. The Plant Cell 15, 2503–2513.

Kim, D.Y., Scalf, M., Smith, L.M., and Vierstra, R.D. (2013). Advanced proteomic analyses yield a deep catalog of ubiquitylation targets in Arabidopsis. The Plant cell 25, 1523–1540.

Kim, S.J., Kim, M.R., Bedgar, D.L., Moinuddin, S.G., Cardenas, C.L., Davin, L.B., Kang, C., and Lewis, N.G. (2004). Functional reclassification of the putative cinnamyl alcohol dehydrogenase multigene family in Arabidopsis. Proc Natl Acad Sci U S A 101, 1455–1460.

Kleinboelting, N., Huep, G., Kloetgen, A., Viehoever, P., and Weisshaar, B. (2012). GABI-Kat SimpleSearch: new features of the Arabidopsis thaliana T-DNA mutant database. Nucleic Acids Res 40, D1211–1215.

Komander, D., and Rape, M. (2012). The ubiquitin code. Annu Rev Biochem 81, 203–229.

Lachaud, C., Prigent, E., Thuleau, P., Grat, S., Da Silva, D., Brière, C., Mazars, C., and Cotelle, V. (2013). 14-3-3-Regulated Ca 2+-dependent protein kinase CPK3 is required for sphingolipid-induced cell death in Arabidopsis. Cell Death & Differentiation 20, 209–217.

Le, M.H., Cao, Y., Zhang, X.C., and Stacey, G. (2014). LIK1, a CERK1-interacting kinase, regulates plant immune responses in Arabidopsis. PLoS One 9, e102245.

Lee, B.-J., Kwon, S.J., Kim, S.-K., Kim, K.-J., Park, C.-J., Kim, Y.-J., Park, O.K., and Paek, K.-H. (2006). Functional study of hot pepper 26S proteasome subunit RPN7 induced by Tobacco mosaic virus from nuclear proteome analysis. Biochemical and biophysical research communications 351, 405–411.

Li, L., Habring, A., Wang, K., and Weigel, D. (2020). Atypical Resistance Protein RPW8/HR Triggers Oligomerization of the NLR Immune Receptor RPP7 and Autoimmunity. Cell Host Microbe 27, 405–417 e406.

Li, Y., Kabbage, M., Liu, W., and Dickman, M.B. (2016). Aspartyl protease-mediated cleavage of BAG6 is necessary for autophagy and fungal resistance in plants. The Plant Cell 28, 233–247.

Liang, X., Ding, P., Lian, K., Wang, J., Ma, M., Li, L., Li, L., Li, M., Zhang, X., Chen, S., et al. (2016). Arabidopsis heterotrimeric G proteins regulate immunity by directly coupling to the FLS2 receptor. Elife 5, e13568.

Liao, D., Cao, Y., Sun, X., Espinoza, C., Nguyen, C.T., Liang, Y., and Stacey, G. (2017). Arabidopsis E3 ubiquitin ligase PLANT U-BOX13 (PUB13) regulates chitin receptor LYSIN MOTIF RECEPTOR KINASE5 (LYK5) protein abundance. The New phytologist 214, 1646–1656.

Liebrand, T.W., van den Burg, H.A., and Joosten, M.H. (2014). Two for all: receptor-associated kinases SOBIR1 and BAK1. Trends in plant science 19, 123–132.

Liu, J., Ding, P., Sun, T., Nitta, Y., Dong, O., Huang, X., Yang, W., Li, X., Botella, J.R., and Zhang, Y. (2013). Heterotrimeric G proteins serve as a converging point in plant defense signaling activated by multiple receptor-like kinases. Plant physiology 161, 2146–2158.

Liu, J., Li, H., Miao, M., Tang, X., Giovannoni, J., Xiao, F., and Liu, Y. (2012a). The tomato UV- damaged DNA-binding protein-1 (DDB1) is implicated in pathogenesis-related (PR) gene expression and resistance to Agrobacterium tumefaciens. Molecular plant pathology 13, 123–134.

Liu, J., Li, H., Miao, M., Tang, X., Giovannoni, J., Xiao, F., and Liu, Y.J.M.p.p. (2012b). The tomato UV-damaged DNA-binding protein-1 (DDB1) is implicated in pathogenesis-related (PR) gene expression and resistance to Agrobacterium tumefaciens. 13, 123–134.

Lu, D., Lin, W., Gao, X., Wu, S., Cheng, C., Avila, J., Heese, A., Devarenne, T.P., He, P., and Shan, L. (2011). Direct ubiquitination of pattern recognition receptor FLS2 attenuates plant innate immunity. Science 332, 1439–1442.

Ma, X., Claus, L.A.N., Leslie, M.E., Tao, K., Wu, Z., Liu, J., Yu, X., Li, B., Zhou, J., Savatin, D.V., et al. (2020). Ligand-induced monoubiquitination of BIK1 regulates plant immunity. Nature 581, 199–203.

Maor, R., Jones, A., Nuhse, T.S., Studholme, D.J., Peck, S.C., and Shirasu, K. (2007). Multidimensional protein identification technology (MudPIT) analysis of ubiquitinated proteins in plants. Mol Cell Proteomics 6, 601–610.

Marino, D., Peeters, N., and Rivas, S. (2012). Ubiquitination during plant immune signaling. Plant physiology 160, 15–27.

Marshall, R.S., Li, F., Gemperline, D.C., Book, A.J., and Vierstra, R.D. (2015). Autophagic Degradation of the 26S Proteasome Is Mediated by the Dual ATG8/Ubiquitin Receptor RPN10 in Arabidopsis. Molecular cell 58, 1053–1066.

Marshall, R.S., and Vierstra, R.D. (2018). Autophagy: The Master of Bulk and Selective Recycling. Annual review of plant biology 69, 173–208.

Meng, X., and Zhang, S. (2013). MAPK cascades in plant disease resistance signaling. Annual review of phytopathology 51, 245–266.

Meyer, D., Pajonk, S., Micali, C., O’Connell, R., and Schulze-Lefert, P.J.T.P.J. (2009). Extracellular transport and integration of plant secretory proteins into pathogen-induced cell wall compartments. 57, 986–999.

Miao, Y., and Zentgraf, U. (2010). A HECT E3 ubiquitin ligase negatively regulates Arabidopsis leaf senescence through degradation of the transcription factor WRKY53. The Plant Journal 63, 179–188.

Monaghan, J., and Li, X. (2010). The HEAT repeat protein ILITYHIA is required for plant immunity. Plant and cell physiology 51, 742–753.

Monaghan, J., Li, X.J.P., and physiology, c. (2010). The HEAT repeat protein ILITYHIA is required for plant immunity. 51, 742–753.

Monaghan, J., Xu, F., Gao, M., Zhao, Q., Palma, K., Long, C., Chen, S., Zhang, Y., and Li, X. (2009). Two Prp19-like U-box proteins in the MOS4-associated complex play redundant roles in plant innate immunity. PLoS Pathogens 5.

Mondragon-Palomino, M., Stam, R., John-Arputharaj, A., and Dresselhaus, T. (2017). Diversification of defensins and NLRs in Arabidopsis species by different evolutionary mechanisms. BMC Evol Biol 17, 255.

Mural, R.V., Liu, Y., Rosebrock, T.R., Brady, J.J., Hamera, S., Connor, R.A., Martin, G.B., and Zeng, L. (2013). The tomato Fni3 lysine-63-specific ubiquitin-conjugating enzyme and suv ubiquitin E2 variant positively regulate plant immunity. Plant Cell 25, 3615–3631.

Murata, S., Yashiroda, H., and Tanaka, K. (2009). Molecular mechanisms of proteasome assembly. Nature reviews Molecular cell biology 10, 104–115.

Nakashima, A., Chen, L., Thao, N.P., Fujiwara, M., Wong, H.L., Kuwano, M., Umemura, K., Shirasu, K., Kawasaki, T., and Shimamoto, K. (2008). RACK1 functions in rice innate immunity by interacting with the Rac1 immune complex. Plant Cell 20, 2265–2279.

Nicaise, V., Joe, A., Jeong, B., Korneli, C., Boutrot, F., Wested, I., Staiger, D., Alfano, J., and Zipfel, C. (2013). Pseudomonas HopU1 affects interaction of plant immune receptor mRNAs to the RNA-binding protein GRP7. EMBO J 32, 701–712.

Okumoto, K., Misono, S., Miyata, N., Matsumoto, Y., Mukai, S., and Fujiki, Y. (2011). Cysteine ubiquitination of PTS1 receptor Pex5p regulates Pex5p recycling. Traffic 12, 1067–1083.

Ordureau, A., Paulo, J.A., Zhang, J., An, H., Swatek, K.N., Cannon, J.R., Wan, Q., Komander, D., and Harper, J.W. (2020). Global Landscape and Dynamics of Parkin and USP30-Dependent Ubiquitylomes in iNeurons during Mitophagic Signaling. Molecular cell 77, 1124–1142 e1110.

Paez Valencia, J., Goodman, K., and Otegui, M.S. (2016). Endocytosis and Endosomal Trafficking in Plants. Annual review of plant biology 67, 309–335.

Palm, D., Streit, D., Shanmugam, T., Weis, B.L., Ruprecht, M., Simm, S., and Schleiff, E. (2019). Plant-specific ribosome biogenesis factors in Arabidopsis thaliana with essential function in rRNA processing. Nucleic Acids Res 47, 1880–1895.

Parker, J.E., Holub, E.B., Frost, L.N., Falk, A., Gunn, N.D., and Daniels, M.J. (1996). Characterization of eds1, a mutation in Arabidopsis suppressing resistance to Peronospora parasitica specified by several different RPP genes. The Plant Cell 8, 2033–2046.

Pastorczyk, M., and Bednarek, P. (2016). The function of glucosinolates and related metabolites in plant innate immunity. In Advances in Botanical Research (Elsevier), pp. 171–198.

Pecher, P., Eschen-Lippold, L., Herklotz, S., Kuhle, K., Naumann, K., Bethke, G., Uhrig, J., Weyhe, M., Scheel, D., and Lee, J. (2014). The A rabidopsis thaliana mitogen-activated protein kinases MPK 3 and MPK 6 target a subclass of ‘VQ-motif’-containing proteins to regulate immune responses. New Phytologist 203, 592–606.

Peng, J., Schwartz, D., Elias, J.E., Thoreen, C.C., Cheng, D., Marsischky, G., Roelofs, J., Finley, D., and Gygi, S.P. (2003). A proteomics approach to understanding protein ubiquitination. Nat Biotechnol 21, 921–926.

Perraki, A., Gronnier, J., Gouguet, P., Boudsocq, M., Deroubaix, A.-F., Simon, V., German-Retana, S., Zipfel, C., Bayer, E., and Mongrand, S. (2017). The plant calcium-dependent protein kinase CPK3 phosphorylates REM1. 3 to restrict viral infection. BioRxiv, 205765.

Pitorre, D., Llauro, C., Jobet, E., Guilleminot, J., Brizard, J.-P., Delseny, M., and Lasserre, E. (2010). RLK7, a leucine-rich repeat receptor-like kinase, is required for proper germination speed and tolerance to oxidative stress in Arabidopsis thaliana. Planta 232, 1339–1353.

Qin, L., Zhou, Z., Li, Q., Zhai, C., Liu, L., Quilichini, T.D., Gao, P., Kessler, S.A., Jaillais, Y., Datla, R., et al. (2020). Specific Recruitment of Phosphoinositide Species to the Plant-Pathogen Interfacial Membrane Underlies Arabidopsis Susceptibility to Fungal Infection. Plant Cell 32, 1665–1688.

Raasi, S., Orlov, I., Fleming, K.G., and Pickart, C.M. (2004). Binding of polyubiquitin chains to ubiquitin-associated (UBA) domains of HHR23A. Journal of molecular biology 341, 1367–1379.

Rayapuram, N., Bigeard, J., Alhoraibi, H., Bonhomme, L., Hesse, A.M., Vinh, J., Hirt, H., and Pflieger, D. (2018). Quantitative Phosphoproteomic Analysis Reveals Shared and Specific Targets of Arabidopsis Mitogen-Activated Protein Kinases (MAPKs) MPK3, MPK4, and MPK6. Mol Cell Proteomics 17, 61–80.

Revers, F., Guiraud, T., Houvenaghel, M.-C., Mauduit, T., Le Gall, O., and Candresse, T. (2003). Multiple resistance phenotypes to Lettuce mosaic virus among Arabidopsis thaliana accessions. Molecular plant-microbe interactions 16, 608–616.

Reyes, F.C., Buono, R., and Otegui, M.S. (2011). Plant endosomal trafficking pathways. Current opinion in plant biology 14, 666–673.

Rodrigues, O., Reshetnyak, G., Grondin, A., Saijo, Y., Leonhardt, N., Maurel, C., and Verdoucq, L. (2017). Aquaporins facilitate hydrogen peroxide entry into guard cells to mediate ABA-and pathogen-triggered stomatal closure. Proceedings of the National Academy of Sciences 114, 9200–9205.

Romero-Barrios, N., and Vert, G. (2018). Proteasome-independent functions of lysine-63 polyubiquitination in plants. The New phytologist 217, 995–1011.

Saracco, S.A., Hansson, M., Scalf, M., Walker, J.M., Smith, L.M., and Vierstra, R.D. (2009). Tandem affinity purification and mass spectrometric analysis of ubiquitylated proteins in Arabidopsis. Plant J 59, 344–358.

Shannon, P., Markiel, A., Ozier, O., Baliga, N.S., Wang, J.T., Ramage, D., Amin, N., Schwikowski, B., and Ideker, T.J.G.r. (2003). Cytoscape: a software environment for integrated models of biomolecular interaction networks. 13, 2498–2504.

Shimizu, Y., Okuda-Shimizu, Y., and Hendershot, L.M. (2010). Ubiquitylation of an ERAD substrate occurs on multiple types of amino acids. Molecular cell 40, 917–926.

Skubacz, A., Daszkowska-Golec, A., and Szarejko, L. (2016). The Role and Regulation of ABI5 (ABA-Insensitive 5) in Plant Development, Abiotic Stress Responses and Phytohormone Crosstalk. Front Plant Sci 7.

Smalle, J., and Vierstra, R.D. (2004). The ubiquitin 26S proteasome proteolytic pathway. Annual review of plant biology 55, 555–590.

Speth, E.B., Imboden, L., Hauck, P., and He, S.Y. (2009a). Subcellular localization and functional analysis of the Arabidopsis GTPase RabE. Plant physiology 149, 1824–1837.

Speth, E.B., Imboden, L., Hauck, P., and He, S.Y. (2009b). Subcellular Localization and Functional Analysis of the Arabidopsis GTPase RabE. 149, 1824–1837.

Spoel, S.H., and Dong, X. (2012). How do plants achieve immunity? Defence without specialized immune cells. Nature reviews Immunology 12, 89–100.

Stegmann, M., Anderson, R.G., Ichimura, K., Pecenkova, T., Reuter, P., Zarsky, V., McDowell, J.M., Shirasu, K., and Trujillo, M. (2012). The ubiquitin ligase PUB22 targets a subunit of the exocyst complex required for PAMP-triggered responses in Arabidopsis. Plant Cell 24, 4703–4716.

Stringlis, I.A., Yu, K., Feussner, K., de Jonge, R., Van Bentum, S., Van Verk, M.C., Berendsen, R.L., Bakker, P.A., Feussner, I., and Pieterse, C.M. (2018). MYB72-dependent coumarin exudation shapes root microbiome assembly to promote plant health. Proceedings of the National Academy of Sciences 115, E5213–E5222.

Su, J., Xu, J., and Zhang, S. (2015). RACK1, scaffolding a heterotrimeric G protein and a MAPK cascade. Trends Plant Sci 20, 405–407.

Su, J., Yang, L., Zhu, Q., Wu, H., He, Y., Liu, Y., Xu, J., Jiang, D., and Zhang, S. (2018). Active photosynthetic inhibition mediated by MPK3/MPK6 is critical to effector-triggered immunity. PLoS biology 16, e2004122.

Szklarczyk, D., Franceschini, A., Wyder, S., Forslund, K., Heller, D., Huerta-Cepas, J., Simonovic, M., Roth, A., Santos, A., Tsafou, K.P., et al. (2015). STRING v10: protein-protein interaction networks, integrated over the tree of life. Nucleic Acids Res 43, D447–452.

Tan, X., Meyers, B.C., Kozik, A., West, M.A., Morgante, M., St Clair, D.A., Bent, A.F., and Michelmore, R.W. (2007). Global expression analysis of nucleotide binding site-leucine rich repeat-encoding and related genes in Arabidopsis. BMC Plant Biol 7, 56.

Tarutani, Y., Morimoto, T., Sasaki, A., Yasuda, M., Nakashita, H., Yoshida, S., Yamaguchi, I., and Suzuki, Y. (2004). Molecular characterization of two highly homologous receptor-like kinase genes, RLK902 and RKL1, in Arabidopsis thaliana. Biosci Biotechnol Biochem 68, 1935–1941.

Thordal-Christensen, H. (2003). Fresh insights into processes of nonhost resistance. Curr Opin Plant Biol 6, 351–357.

Tronchet, M., Balague, C., Kroj, T., Jouanin, L., and Roby, D. (2010). Cinnamyl alcohol dehydrogenases-C and D, key enzymes in lignin biosynthesis, play an essential role in disease resistance in Arabidopsis. Molecular plant pathology 11, 83–92.

Trujillo, M., and Shirasu, K. (2010). Ubiquitination in plant immunity. Current opinion in plant biology 13, 402–408.

Udeshi, N.D., Svinkina, T., Mertins, P., Kuhn, E., Mani, D.R., Qiao, J.W., and Carr, S.A. (2013). Refined Preparation and Use of Anti-diglycine Remnant (K-epsilon-GG) Antibody Enables Routine Quantification of 10,000s of Ubiquitination Sites in Single Proteomics Experiments. Molecular & Cellular Proteomics 12, 825–831.

Urano, D., Chen, J.G., Botella, J.R., and Jones, A.M. (2013). Heterotrimeric G protein signalling in the plant kingdom. Open Biol 3, 120186.

Üstün, S., Sheikh, A., Gimenez-Ibanez, S., Jones, A., Ntoukakis, V., and Börnke, F. (2016). The proteasome acts as a hub for plant immunity and is targeted by Pseudomonas type III effectors. Plant Physiology 172, 1941–1958.

van Nocker, S., Walker, J.M., and Vierstra, R.D. (1996). The Arabidopsis thaliana UBC7/13/14 genes encode a family of multiubiquitin chain-forming E2 enzymes. The Journal of biological chemistry 271, 12150–12158.

Vierstra, R.D. (2009). The ubiquitin-26S proteasome system at the nexus of plant biology. Nat Rev Mol Cell Biol 10, 385–397.

Wang, J., Grubb, L.E., Wang, J., Liang, X., Li, L., Gao, C., Ma, M., Feng, F., Li, M., Li, L., et al. (2018). A Regulatory Module Controlling Homeostasis of a Plant Immune Kinase. Molecular cell.

Wang, L., Wen, R., Wang, J., Xiang, D., Wang, Q., Zang, Y., Wang, Z., Huang, S., Li, X., and Datla, R.J.N.P. (2019). Arabidopsis UBC 13 differentially regulates two programmed cell death pathways in responses to pathogen and low-temperature stress. 221, 919–934.

Wawrzynska, A., Rodibaugh, N.L., and Innes, R.W. (2010). Synergistic activation of defense responses in Arabidopsis by simultaneous loss of the GSL5 callose synthase and the EDR1 protein kinase. Mol Plant Microbe Interact 23, 578–584.

Wen, R., Torres-Acosta, J.A., Pastushok, L., Lai, X., Pelzer, L., Wang, H., and Xiao, W. (2008). Arabidopsis UEV1D promotes Lysine-63-linked polyubiquitination and is involved in DNA damage response. The Plant cell 20, 213–227.

Yao, C., Wu, Y., Nie, H., and Tang, D. (2012). RPN1a, a 26S proteasome subunit, is required for innate immunity in Arabidopsis. The Plant journal : for cell and molecular biology 71, 1015–1028.

Yu, X., Feng, B., He, P., and Shan, L. (2017). From Chaos to Harmony: Responses and Signaling upon Microbial Pattern Recognition. Annual review of phytopathology 55, 109–137.

Zamioudis, C., Hanson, J., and Pieterse, C.M. (2014). β-Glucosidase BGLU 42 is a MYB 72- dependent key regulator of rhizobacteria-induced systemic resistance and modulates iron deficiency responses in A rabidopsis roots. New Phytologist 204, 368–379.

Zhao, J., Zhou, H., Zhang, M., Gao, Y., Li, L., Gao, Y., Li, M., Yang, Y., Guo, Y., and Li, X. (2016). Ubiquitin-specific protease 24 negatively regulates abscisic acid signalling in Arabidopsis thaliana. Plant, cell & environment 39, 427–440.

Zhao, Y., Wu, G., Shi, H., and Tang, D. (2019). RECEPTOR-LIKE KINASE 902 Associates with and Phosphorylates BRASSINOSTEROID-SIGNALING KINASE1 to Regulate Plant Immunity. Mol Plant 12, 59–70.

Zhou, B., and Zeng, L. (2017). Conventional and unconventional ubiquitination in plant immunity. Molecular plant pathology 18, 1313–1330.

Zhou, J.-M., and Zhang, Y. (2020). Plant Immunity: Danger Perception and Signaling. Cell.

Zhou, J., He, P., and Shan, L. (2014). Ubiquitination of plant immune receptors. Methods Mol Biol 1209, 219–231.

Zhou, J., Liu, D., Wang, P., Ma, X., Lin, W., Chen, S., Mishev, K., Lu, D., Kumar, R., Vanhoutte, I., et al. (2018). Regulation of Arabidopsis brassinosteroid receptor BRI1 endocytosis and degradation by plant U-box PUB12/PUB13-mediated ubiquitination. Proc Natl Acad Sci U S A 115, E1906–E1915.

Zhou, J., Lu, D., Xu, G., Finlayson, S.A., He, P., and Shan, L. (2015). The dominant negative ARM domain uncovers multiple functions of PUB13 in Arabidopsis immunity, flowering, and senescence. J Exp Bot 66, 3353–3366.

